# Evaluating exon-skipping therapies targeting the central nervous system in Duchenne muscular dystrophy using high-resolution spatial transcriptomics

**DOI:** 10.1101/2025.10.20.683239

**Authors:** Qirong Mao, Alireza Ahmadi, Sharon de Vries, Laura G.M. Heezen, Ophélie Vacca, Mathilde Doisy, Annemieke Aartsma-Rus, Maaike van Putten, Aurélie Goyenvalle, Ahmed Mahfouz, Pietro Spitali

## Abstract

Duchenne muscular dystrophy (DMD) is marked by progressive muscle degeneration due to dystrophin deficiency. Despite cognitive impairments in up to one-third of patients, central nervous system (CNS)-targeted dystrophin restoration remains relatively underexplored compared to muscle-focused therapies. Since dystrophin is expressed in the brain during development and postnatally, it is important to assess which aspects of CNS pathology could be rescued. However, studying *Dmd* is difficult due to its size, low-abundance transcripts, and with multiple isoforms, limiting the sensitivity and isoform resolution of standard sequencing methods. To overcome these limitations, we applied Xenium spatial transcriptomics to target splice junctions, enabling isoform-specific and exon 51 skipping events detection across brain regions and cell types. Using this approach, we analyzed *mdx52* mice lacking the full-length and Dp140 isoforms, treated with two exon 51 skipping therapies (antisense oligonucleotides or AAV-U7ex51). We observed distinct spatial expression patterns in the wild type brain between the full-length isoforms (Dp427c/m/p1) and shorter isoforms (Dp71 and Dp40). The full-length isoforms were predominantly expressed in cortex layers 2/3–6b and the CA1 region, while the shorter isoforms were localized to cortex layer 1 and the dentate gyrus. Among the exon skipping therapies tested, U7ex51 delivered neonatally induced broad exon skipping in neurons and effectively restored full-length isoforms in the targeted cells. This study introduces the first subcellular-resolution spatial transcriptomic atlas of dystrophin presence and rescue in the CNS and demonstrates a framework for evaluating gene therapies in spatially and transcriptionally complex tissues such as the brain.

## INTRODUCTION

Duchenne muscular dystrophy (DMD) is an X-linked neuromuscular disorder caused by mutations in the *DMD* gene, which encodes dystrophin. Nonsense and frameshift mutations in the *DMD* gene cause the absence of functional dystrophin in DMD. Along with the progressive loss of muscle tissue and functionality, DMD is also associated with cognitive disorders. Nearly one-third of DMD patients show mild to severe mental retardation with Full-Scale Intelligence Quotient (FSIQ) lower than 70 and overall mean FSIQ decreasing by one standard deviation compared to normal populations (*1–3*). Additionally, a higher prevalence of neurodevelopmental disorders was observed among DMD patients, such as emotional dysregulation, epilepsy, attention-deficit/hyperactivity disorder (ADHD), autism spectrum disorder (ASD), and obsessive-compulsive disorders (*4–9*). Recent studies have also shown that DMD patients frequently exhibit academic challenges, such as reading difficulties and deficits in arithmetic abilities (*10–12*). In the last decades, developments in care strategies have improved disease progression significantly. However, relatively little attention has been given to investigating brain comorbidities in DMD. These challenges continue to negatively impact the quality of life of DMD patients, underscoring the urgent need for further research and targeted interventions.

The *DMD* gene has seven tissue-specific promoters, enabling the production of various dystrophin protein (Dp) isoforms named by their molecular size. Three full-length 427 kDa isoforms are encoded by three separate promoters located upstream of the first exon: Dp427m is mainly expressed in skeletal, cardiac and smooth muscle, Dp427c is present in neurons of the cerebral cortex and cerebellum, and Dp427p is expressed in the Purkinje cells in the cerebellum (*13, 14*). In addition, shorter dystrophin isoforms (Dp260, Dp140, Dp116, Dp71, and Dp40) are transcribed by downstream promoters located in the introns of the *DMD* gene. Dp260 is predominantly localized to the retina, and Dp116 is primarily expressed in the peripheral nerve (*15, 16*). ‘Dp140, Dp71, and Dp40 are expressed in the CNS as well as in other tissues. Dp140 is also expressed in the kidney, whereas Dp71 and Dp40 are broadly expressed across multiple tissues and are highly expressed in the CNS (*13, 17*).

Brain comorbidities have been linked to the mutation site along the gene, with severity and frequency of phenotypes being higher for patients carrying pathogenic variants located in the distal part of the *DMD* gene, leading to a cumulative loss of dystrophin isoforms. Meta analysis has shown that DMD patients lacking Dp427 have lower average IQ scores, with progressive reduction when Dp140 and Dp71 are also absent (*18*). Distal mutations affecting Dp140/Dp71 are associated with a higher prevalence of ASD and ADHD diagnoses, as well as emotional and behavioral problems, although variability between patients remains substantial (*19–21*). In addition, DMD patients missing only Dp427 show better motor performance than those also missing Dp140 and/or Dp71 (*22, 23*).

Over the past decades, dystrophin restoring therapies such as antisense oligonucleotide (ASO) mediated exon skipping have shown promise in DMD. ASOs are small synthetic molecules that interfere with pre-mRNA splicing, thereby causing the exclusion of specific exon(s) from the mature mRNA. Such an approach has been shown to restore the open reading frame and to allow the synthesis of truncated but partially functional dystrophins (*24*). Clinical trials have demonstrated only minimal dystrophin restoration in the skeletal muscle of patients with DMD following ASO treatment, leading to the accelerated approval by the U.S. Food and Drug Administration (FDA) of four ASOs and by Japan’s Ministry of Health, Labour and Welfare (MHLW) of one ASO (*25–28*).

In addition to ASOs, antisense sequences can also be introduced in cells through viral delivery of the antisense sequence as part of a uridine-rich 7 small nuclear RNA (U7snRNA) cassette. U7snRNA is a part of the small nuclear ribonucleoprotein and participates in 3’ end processing of histone pre-mRNA (*29*). With the modification in the binding site for Sm/Lsm protein, U7snRNA can mediate splicing and induce skipping/inclusion of target exons (30). Delivery of U7snRNA with an antisense sequence has been explored using adeno-associated viral vectors (AAV), showing therapeutic potential in DMD mouse and dog models, and has been used in a small-scale open label clinical trial (*31–33*).

Despite therapeutic advances in dystrophin restoration targeting muscle in DMD patients, the brain is left untreated as these ASOs cannot cross the blood-brain-barrier. Efforts to develop exon skipping treatment addressing brain comorbidities in DMD remain scarce and in their infancy. One of the barriers is the limited knowledge of dystrophin isoforms’ function and localization across brain regions and cell types in the brain (*14, 34*).

Preliminary studies explored the potential of ASOs to restore dystrophin expression and alleviate brain-related symptoms in *mdx52* mice (which carry a deletion of exon 52 and consequently lack expression of Dp427 and Dp140) (*35*). A single intracerebroventricular (ICV) administration of ASOs targeting *Dmd* exon 51 achieved partial restoration of Dp427 (5-15%) in brain regions expressing dystrophins, including the hippocampus, cerebellum, and cortex. Behaviorally, this treatment significantly reduced anxiety and unconditioned fear but only partially improved fear memory, suggesting that the modest behavioral benefits could result from the limited levels of Dp427 restoration and/or the relatively late (adult) timing of treatment. These findings raised the key question of whether higher dystrophin restoration or earlier intervention might lead to stronger functional recovery. This motivated us to explore alternative delivery strategies. In the present study, we therefore evaluated ICV delivery of AAV-U7 in neonatal *mdx52* mice, directly comparing its effects with the previously described ICV delivery of ASO in adult animals (*35*).

In this research, we applied Xenium spatial transcriptomics to map dystrophin isoforms localization and exon skipping events in the murine brain. This approach allows us to describe the spatial distribution of dystrophin isoforms in the brain, at cellular level. We further evaluated the therapeutic effects of two different exon skipping treatments in the brain of the *mdx52* mouse model, lacking full length and Dp140 isoforms.

## RESULTS

### Quality assessment and annotation of spatial transcriptomics data

We performed Xenium *in situ* sequencing (10X Genomics) on hemi-coronal brain sections from 12 individual mice across 4 groups: wildtype (N=3), *mdx52* mice treated with PBS (N=3), *mdx52* mice treated with either ASO (tcDNA) (N=3) or AAV-U7snRNA targeting *Dmd* exon 51 skipping (U7ex51) (N=3) (Material and Method). All samples, including those from WT, were collected at 14 weeks of age (Fig. 1A).

**Fig. 1.**
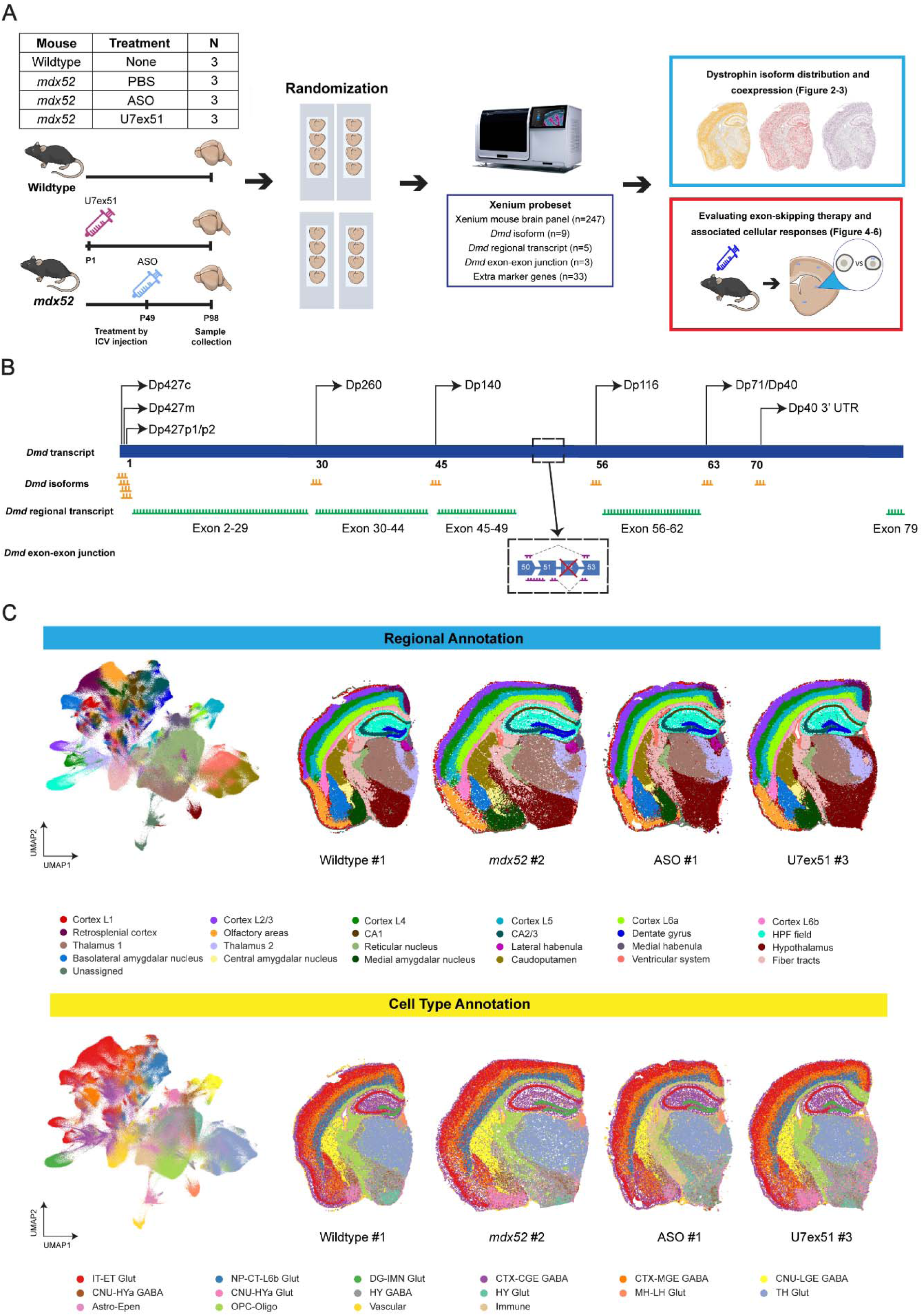
Spatial profiling of brain structure and cellular composition in DMD and wildtype mouse models. **(A)** Overview of the experimental design and downstream data analysis workflow. **(B)** Schematic representation of the custom Xenium panel design, including probes targeting isoforms promotor or untranslated regions (probes shown in yellow), *Dmd* regional transcripts (probes shown in green), and exon–exon junctions (probes shown in purple). **(C)** Uniform Manifold Approximation and Projection (UMAP) plots for all samples, with spatial plots showing (Top) regional and (Bottom) cell type annotation of selected replicates per group. (Astro, astrocyte; CGE, caudal ganglionic eminence; CNU, cerebral nuclei; CT, corticothalamic; CTX, cerebral cortex; Epen, ependymal; ET, extratelencephalic; GABA, GABAergic; Glut, glutamatergic; HPF, hippocampal formation; HY, hypothalamus; HYa, anterior hypothalamic; IMN, immature neurons; IT, intratelencephalic; L6b, layer 6b; LGE, lateral ganglionic eminence; LH, lateral habenula; MGE, medial ganglionic eminence; MH, medial habenula; NP, near-projecting; Oligo, oligodendrocytes; OPC, oligodendrocyte precursor cells; TH, thalamus, named by (*36*)).

Here, we employed the pre-designed Xenium mouse brain panel with 247 genes, in addition to a custom add-on panel of 50 genes (Data file S1). To investigate the spatial distribution of dystrophin isoforms across brain regions and cell types, we included custom probes to detect *Dmd* isoforms and *Dmd* transcript regions (proximal, middle, and distal). We also included *Dmd* exon-exon junction probes to evaluate exon 51 skipping therapeutic efficacy (Fig. 1B).

Overall, we detected 949,583 cells with an average of 91 (91.34 ± 31.54) genes per cell across the 12 samples (Fig. S1-3). Based on the matching of the marker genes in our panel with the single-cell RNA-seq atlas of the mouse brain from the Allen Brain Atlas (*36*), we annotated segmented cells at both the regional and cell type level, identifying 24 brain regions and 16 distinct cell types (Fig. 1C; Fig. S4-6).

### Spatial localization of dystrophin isoforms in the wild type brain

We examined the spatial distribution of multiple dystrophin isoforms across our samples using promoter specific probes. Probes for the Dp260 and Dp140 promoters were excluded from the downstream analysis due to insufficient signal (Fig. S7). Dp427c was the predominant isoform in the WT brain, while Dp427m, Dp427p1, the first exon shared by Dp71 and Dp40 (Dp71 + Dp40), and the 3’UTR of Dp40 (Dp40) had lower expression levels. Dp427p2 showed the lowest expression level across the brain (Fig. 2A, B).

**Fig. 2.**
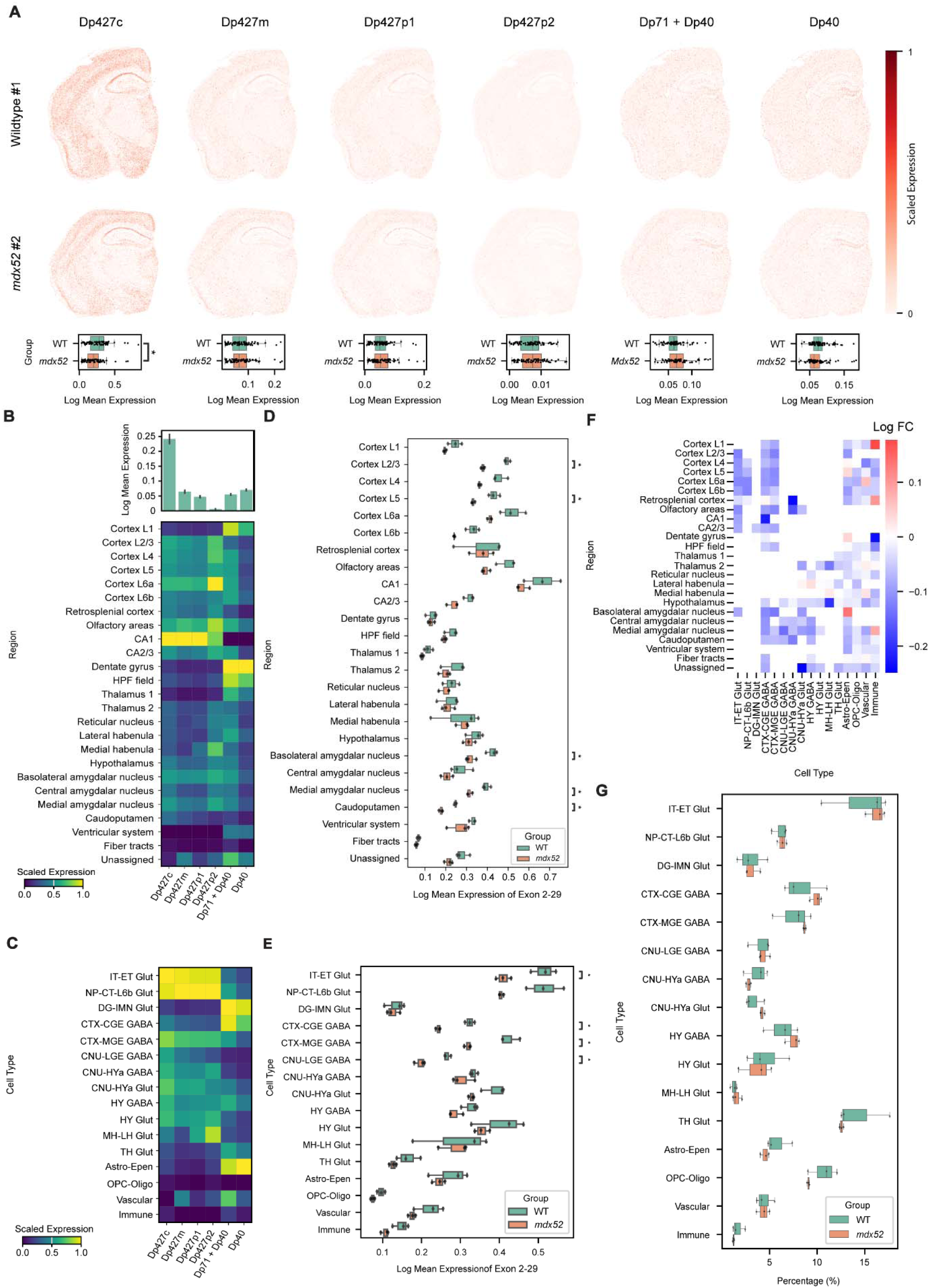
Distinct distribution patterns of dystrophin isoforms and the differential expression of the full-length *Dmd* isoform across brain regions and cell types between wildtype and *mdx52* mice. **(A)** (Top) Spatial distribution of dystrophin isoforms in WT and *mdx52* mouse brains colored by scale expression of each *Dmd* isoform. (Bottom) Box plot with overlaid dotplot showing the log-transformed mean expression of each *Dmd* isoform across brain regions of WT and *mdx52* mice. Statistical analysis by the linear regression test with Benjamini–Hochberg (BH) correction **(B-C)** Expression of dystrophin isoforms in WT mice and their log-normalized expression across brain regions and cell types. Results are shown in mean ± Standard Error of the Mean (SEM). **(D-E)** Expression of exon 2-29 across brain regions and cell types in WT and *mdx52* mice. Statistical analysis by the one-sided t-test with BH correction. **(F)** Log fold changes in exon 2-29 expression across brain regions and cell types between WT and *mdx52* mice. **(G)** Cell type proportions in brain samples of WT and *mdx52* mice. \**P* < 0.05.

The expression of dystrophin isoforms across brain regions revealed two common patterns: one for the full-length isoforms (Dp427c, Dp427m, Dp427p1) and one for the shorter isoforms (Dp71 + Dp40, Dp40). The full-length isoforms were mainly expressed in the cortex from layer 2/3 to layer 6b, olfactory areas, CA1, CA2/3, and basolateral amygdalar nucleus, while the shorter isoforms were highly expressed in cortex layer 1, the dentate gyrus, and the HPF field (Fig. 2B). We also found a distinct pattern at the cellular level: the full-length isoforms showed high expression in IT-ET Glut, NP-CT-L6b Glut, CTX-MGE GABA, CNU-HYa GABA, and HY Glut, while the shorter isoforms were predominantly expressed in DG-IMN Glut, CTX-CGE GABA and Astro-Epen cells. Moreover, we observed that Dp427m showed higher expression levels in vascular cells compared to the other full-length isoforms (Fig. 2C).

### *Mdx52* mice show altered full-length isoform expression

To better understand genotype-specific differences in *Dmd* isoform expression, we compared their expression between WT and *mdx52* mice. Dp427c was found to be less expressed in *mdx52* mice (linear regression test; β = −0.10, adjusted *P* = 0.0139) (Fig. 2A), while the other full-length and shorter isoforms did not show statistically significant differences. However, all *Dmd* regional probes, which included a larger number of probe-sets per region than isoform probes, leading to a stronger signal, showed significantly lower expression in *mdx52* mice compared to WT mice (Fig. S8A). We therefore examined the expression of the exon 2–29 probe across brain regions and cell types (Fig. S8B-D).

Across brain regions, we observed that the expression of the full-length isoforms tended to decrease in the *mdx52* group. In the isocortex, the full-length isoforms had significantly lower expression in layer 2/3 and 5, and also displayed reduced expression in layer 4 and 6b, although this did not reach significance (layer 4: adjusted *P* = 0.08; layer 6b: adjusted *P* = 0.09; one-sided t-test). In the hippocampal formation, the full-length isoforms showed a down-regulated trend in the CA2/3 region of *mdx52* mice, but the difference was not statistically significant. For other brain regions, the full-length isoforms showed significantly lower expression in the basolateral and medial amygdalar nucleus, and caudo-putamen in the *mdx52* group (Fig. 2D).

Regarding cell types, we observed that the full-length isoforms were significantly lower expressed in neuronal cells in the *mdx52* brain. This includes IT-ET Glut, CTX-CGE GABA, CTX-MGE GABA, and CNU-LGE GABA cells, which are primarily located in the isocortex, hippocampus, and basolateral amygdalar nucleus. Similar trends were noted in NP-CT-L6b Glut cells located in the cortex L5/6, although this did not reach statistical significance (one-sided t-test; adjusted *P* = 0.06). Among non-neuronal cell types, our data also suggests a trend towards reduced expression in OPC-Oligo cells and vascular cells (OPC-Oligo: adjusted *P* = 0.06; Vascular: adjusted *P* = 0.12; one-sided t-test) (Fig. 2E-F). Interestingly, the observed changes in expression of the full-length isoforms were not explained by differences in cell type proportions between WT and *mdx52* mice (Fig. 2G).

### Spatial divergence and coexpression of full-length and shorter dystrophin isoforms in the wildtype mouse brain

We next sought to identify regions and cell types showing mutually exclusive or coexpression of full-length and short *Dmd* isoforms in the WT mouse brain. For this purpose, we used promoter specific probes as only these could detect the expression of specific short isoforms. We categorized the dystrophin isoforms into two groups: the full-length isoforms (Dp427c/m/p1), excluding isoform Dp427p2 due to its low expression, and shorter isoforms (Dp71+Dp40, Dp40).

Cells expressing only the full-length *Dmd* isoforms were predominantly found among neuronal populations, comprising 65% to 90% of all *Dmd*-positive cells. These included IT-ET Glutamatergic cells, NP-CT-L6b Glutamatergic cells, and CTX-MGE GABAergic cells, which are primarily located in the isocortex, hippocampal CA region, and amygdala. The remainder of the cells expressing only the full-length *Dmd* isoforms consisted of CNU-LGE GABAergic, CNU-HYa GABAergic, and CNU-HYa Glutamatergic populations, distributed across the thalamus, hypothalamus, amygdala, and caudoputamen. Notably, a high proportion of cells expressing only the full-length *Dmd* isoforms was also detected among OPC-Oligo populations (Fig. 3A, D; Fig. S9A, B).

**Fig. 3.**
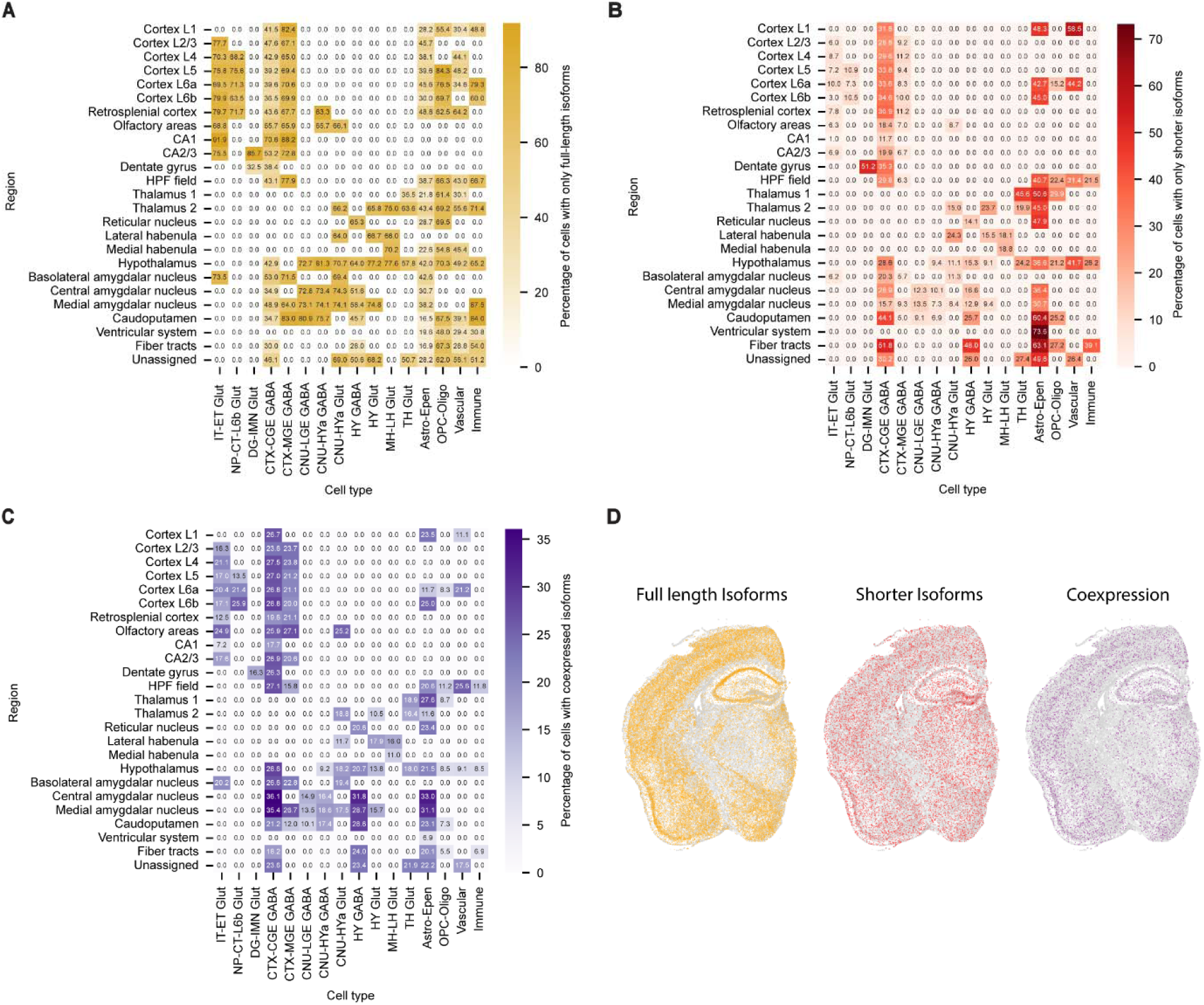
Localization and coexpression of the full-length and shorter *Dmd* isoforms in the wildtype mouse brain. **(A-C)** Heatmap showing the relative expression rate of **(A)** full-length *Dmd* isoforms (Dp427c/m/p1) **(B)** shorter *Dmd* isoforms (Dp71+Dp40, Dp40) **(C)** coexpressed isoforms across the region and cell types in the wildtype mouse brain. **(D)** Spatial distribution of cells solely expressing full-length or shorter *Dmd* isoforms, and cells coexpressing these *Dmd* isoforms in the wildtype mouse brain.

Cells with exclusive expression of the shorter *Dmd* isoforms were found among both neuronal and non-neuronal populations. For instance, approximately 50% of *Dmd* isoform-positive DG-IMN Glut cells in the dentate gyrus and around 30% of CTX-CGE GABA *Dmd* isoform-positive cells across various brain regions expressed only the shorter isoforms. Additionally, our data revealed that astro-ependymal cells also exhibited exclusive expression of shorter isoforms, comprising 35% to 74% of *Dmd* isoform-positive cells within this population (Fig. 3B, D).

Next, we investigated the proportion of cells coexpressing both the full-length and shorter *Dmd* isoforms. These cells were predominantly located in the isocortex, hippocampus, and amygdala. Notably, CTX-CGE GABA neurons showed a notable prevalence of coexpression, representing approximately 25% of the *Dmd*-positive population within this subtype. A similar pattern was also shown in IT-ET Glut and CTX-MGE GABA cells, but with lower coexpression rates (Fig. 3C-D). This suggests that the coexpression between full-length and shorter *Dmd* isoforms is mainly driven by certain cell types (Fig. 3C-D).

### Efficacy of exon 51 skipping events in the brain after therapeutic intervention

To evaluate exon 51 skipping therapies in the DMD brain, *mdx52* mice were treated with two distinct approaches (ASO and U7ex51-mediated exon skipping), and exon 51 skipping was verified using RT-PCR analysis in the isocortex and hippocampus (Fig. 4A). Comparison of exon skipping rates between the two treatments revealed that U7ex51 treatment induced higher exon 51 skipping in both regions compared to ASO treatment, although the differences were not statistically significant (Hippocampus: *P* = 0.2; Isocortex: *P* = 0.2; Mann-Whitney U test). This difference is likely attributable to the distinct therapeutic strategies (U7ex51 versus ASO) and the timing of administration, since U7ex51 was delivered at the neonatal stage whereas ASO treatment was performed in adult mice (6–7 weeks old) (Fig. 4B).

**Fig. 4.**
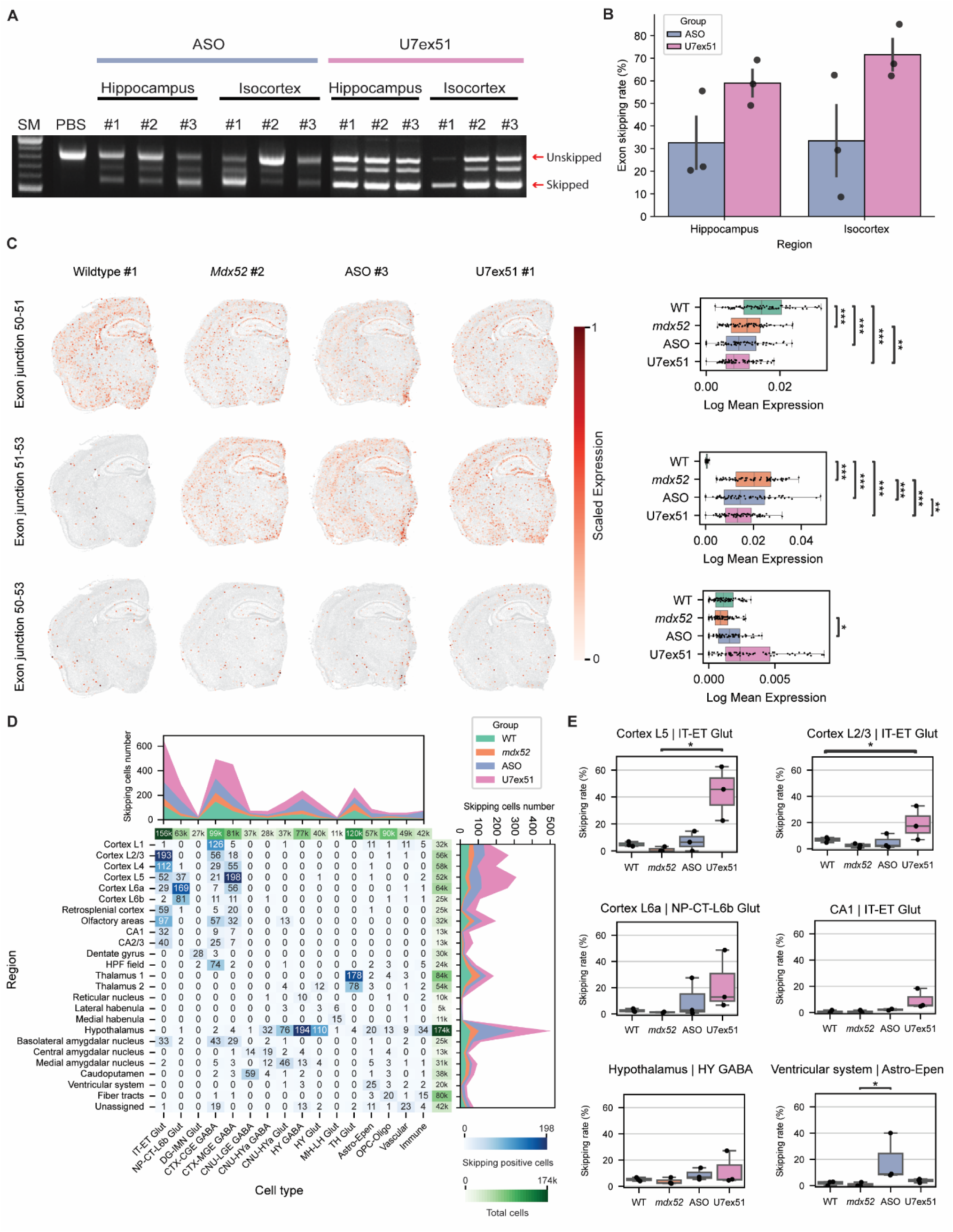
Exon skipping events in the brain induced by exon 51 skipping treatments. **(A)** Quantification of mRNA levels corresponding to exon 52 deletion and combined exon 52 deletion with exon 51 skipping in the hippocampus and isocortex, measured by RT-qPCR across ASO and U7ex51-treated samples. SM: Size Marker **(B)** Box plot with overlaid dot plot showing exon 51 skipping rate in hippocampus and isocortex across ASO and U7ex51 samples measured by RT-qPCR. Results are shown as mean ± SEM. Statistical analysis by the two-sided Mann-Whitney U test. **(C)** (Left) Spatial distribution of cells expressing unskipped transcripts, exon 52 deletion, and combined exon 52 deletion with exon 51 skipping. (Right) Box plot with overlaid dot plot showing the log-transformed mean expression of each transcript type across brain regions per sample across different groups. Dots are displayed only within the interquartile range (Q1–Q3) of each box plot. Statistical analysis by the linear mixed model, and post-hoc pairwise comparisons were performed using estimated marginal means (emmeans) with Tukey’s adjustment for multiple test correction. **(D)** Frequencies of exon 51 skipping events across brain regions and cell types for all samples in each group. **(E)** Box plot showing exon 51 skipping rates per sample in selected brain regions and cell types. Statistical analysis by the Kruskal–Wallis test, followed by Dunn’s post-hoc test with Benjamini–Hochberg correction for multiple comparisons. \**P* < 0.05; \*\**P* < 0.01; \*\*\**P* < 0.001.

To spatially resolve exon 51 skipping events in the brain, we designed exon–exon junction probes targeting the *Dmd* transcript for Xenium: exon 50-51 junction (unskipped transcripts), exon 51-53 junction (exon 52 deleted transcripts), and exon 50-53 junction (skipped transcripts lacking both exons 51 and 52) (Fig. 1C). Our data showed that the expression of the exon 50-51 junction in the WT group is higher than in the *mdx52* groups, confirming reduced expression of *Dmd* transcript in *mdx52* mice. The expression of exon 51-53 junction detecting exon 52 deletion showed higher expression in the *mdx52* group and lower expression in the WT group, confirming the specificity of the probe detecting the deletion of exon 52. For the 50-53 junction probes, the ASO group showed significantly higher expression compared to the *mdx52* group. While the U7ex51 group also exhibited increased expression, this difference was not statistically significant (*P* = 0.06; estimated marginal means analysis). These observations are consistent with the RT-PCR results, supporting the robustness and specificity of the exon junction probes (Fig. 4C).

We next mapped exon 51 skipping events across brain regions and cell types (Fig. 4D) and observed that these occurred predominantly in the neuronal population. Cortical neurons exhibited abundant skipping events, along with TH Glut neurons in the thalamus, and both HY GABA and HY Glut neurons in the hypothalamus. We also observed exon 51 skipping in neuronal cells within the hippocampal CA regions and the amygdala, but these events were less frequent, despite the high expression levels of full-length *Dmd* isoforms typically observed in these brain regions in WT mice (Fig. 2B).

Furthermore, we examined specific brain regions and cell types to compare skipping rates between therapeutic groups (Fig. 4E). Our data showed that both ASO and U7ex51 treated brains had increased skipping rates compared to the untreated *mdx52* brains in cortical neurons, such as IT-ET Glut in cortex L2/3 and L5, NP-CT-L6b Glut in cortex L6, and HY GABA in the hippocampus, with the U7ex51 group exhibiting consistently higher skipping rates than the ASO group. However, we found that the ASO group had a higher skipping rate in astrocyte-ependymal cells in the ventricular system, suggesting that the delivery of ASOs and U7ex51 may be cell-type specific. Surprisingly, despite high Dp427c expression in the IT-ET Glut of the CA1 region, both therapeutic groups exhibited an overall lower skipping rate.

### Full-length isoform restoration in exon 51 skipped cells

Given the successful induction of exon skipping by both therapeutics in our study, we next assessed whether the exon skipping events led to the restoration of full-length dystrophin protein in the hippocampus and isocortex by Western blot (Fig. 5A). We confirmed consistent restoration of the Dp427 isoform in both regions across all treated mice, regardless of the therapeutic approach. The ASO group showed an average of 8.9% ± 2.1% Dp427 protein restoration of WT level in the hippocampus and 7.3% ± 4.6% in the isocortex (mean ± SEM). In comparison, the U7ex51 group showed a higher abundance of Dp427 expression with an average of 41.9% ± 9.6% and 28.0% ± 10.9% in the hippocampus and isocortex, respectively (mean ± SEM; Hippocampus: *P* =0.1; Isocortex: *P* = 0.2; Mann-Whitney U test).

**Fig. 5.**
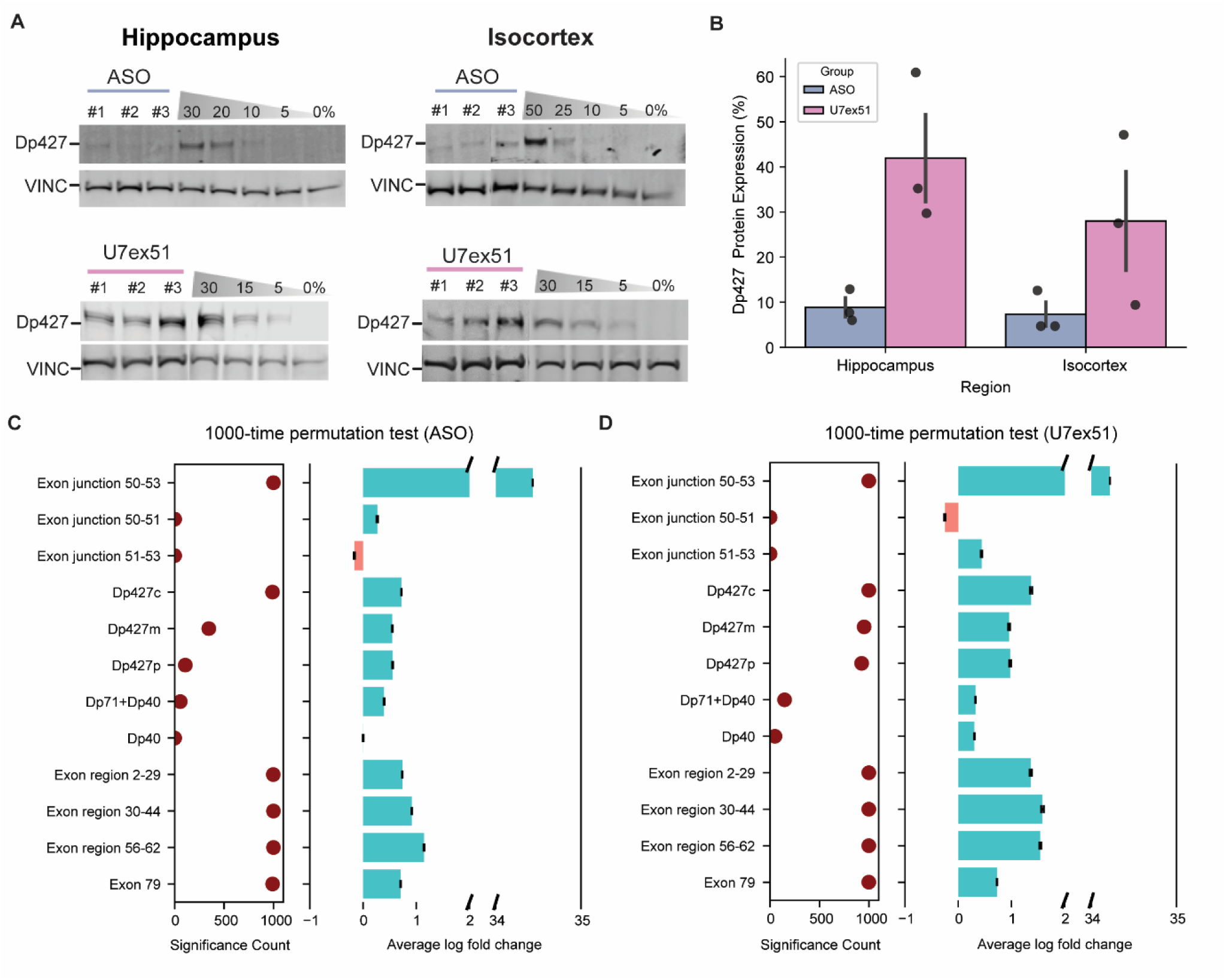
Cellular restoration of *Dmd* isoforms after exon 51 skipping therapy. **(A)** Quantification of Dp427 protein restoration levels by Western blot in ASO- and U7ex51-treated samples. Protein levels were quantified using standard curves generated on separate gels. Each gel included 0% to 50% wildtype lysate dilutions mixed with *mdx52* lysate. VINC; vinculin. **(B)** Percentage of Dp427 isoform protein restoration in the hippocampus and cortex across ASO and U7ex51 treatment groups, measured by Western blot. Results are shown as mean ± SEM. Statistical analysis by the two-sided Mann-Whitney U test. **(C-D)** Differential expression analysis of *Dmd* transcripts between exon 51-skipped and matched unskipped cells in **(C)** ASO-treated and **(D)** U7ex51-treated groups, assessed using the Wilcoxon test with 1000 permutations. The dot plot displays the number of significant outcomes across 1000 permutations, while the bar plot shows the average log fold change ± SEM in exon 51-skipped cells across permutations.

We then assessed whether *Dmd* expression changes were coupled to exon 51 skipping events by comparing the expression of all *Dmd*-related transcripts between exon 51–skipped and non-skipped cells within each therapeutic group. In the 811 cells presenting exon 51 skipping events in the ASO group (representing 0.4% of all cells in this group), we observed elevated expression of the Dp427c isoform and all *Dmd* regional transcripts, indicating restoration of Dp427 expression levels following exon 51 skipping (Fig. 5C). For the 1299 cells presenting exon 51 skipping events in the U7ex51 group (representing 0.6% of all cells in this group), all full-length *Dmd* isoforms (Dp427c, Dp427m, and Dp427p), along with all regional *Dmd* transcripts, showed higher expression, indicating more widespread and robust dystrophin restoration in the U7ex51-treated group (Fig. 5D).

### ASO treatment induces regionally localized microglial activation

When comparing the overall transcriptomics profiles of both exon skipping treatments, one feature that stood out was the difference in immune cell populations (*37*). We further examined immune cell populations to assess whether exon 51 skipping could induce an immune response in the CNS. We first identified immune cell subpopulations by re-clustering all annotated immune cells, resulting in 9 immune subpopulations (Fig. 6A). We noticed an overall expansion of immune cell populations in ASO-treated samples (Fig. 6B). Among these, a specific subcluster (Cluster 1), characterized by the expression of *Trem2*, *Spp1*, and *Cd68* (Fig. 6C), was predominantly enriched in the ASO-treated group (Fig. 6B). The marker gene profile of this cluster suggests an increase in immune cells exhibiting an activated microglial phenotype (*38–40*).

**Fig. 6.**
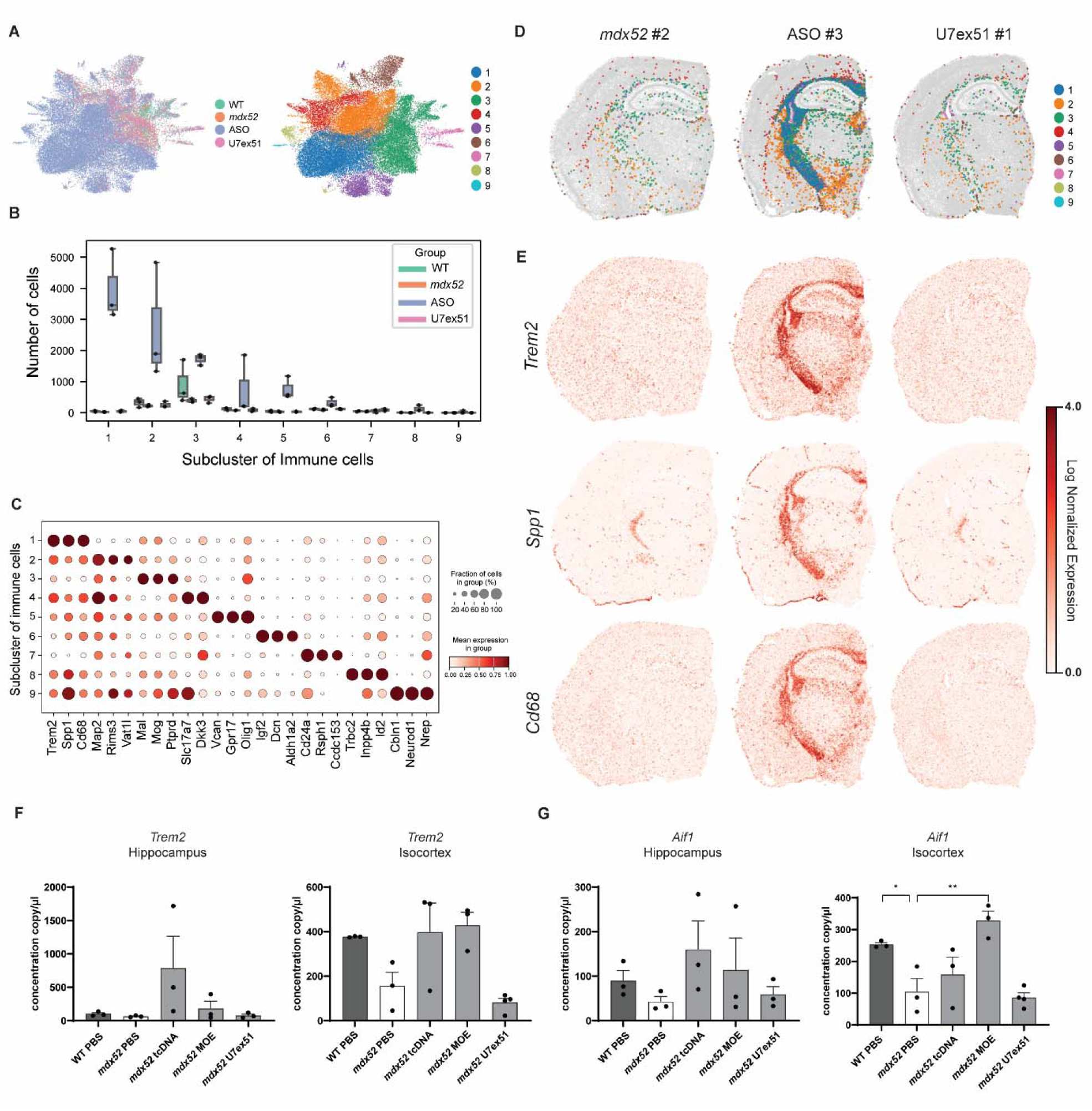
Immune response in ASO-treated brains of *mdx52* mice. **(A)** UMAP visualization of subclusters of immune cells across the treatment groups. **(B)** Box plot with overlaid dot plot showing cell counts for each immune cell subcluster across samples from all groups. **(C)** Dot plot showing marker gene expression profiles for each immune cell subcluster. **(D)** Spatial distribution of immune cell subclusters in tissue samples from *mdx52*, ASO-treated, and U7ex51-treated mice. **(E)** Spatial expression patterns of *Spp1*, *Cd68*, and *Trem2* in samples from *mdx52*, ASO-treated, and U7ex51-treated mice (log-normalized). **(F–G)** Droplet digital PCR (ddPCR) quantification of RNA levels for **(F)** *Trem2* and **(G)** *Aif1* across the hippocampus and isocortex in samples from WT, *mdx52*, ASO-treated (tcDNA and 2′-O-methoxyethylribose [MOE]), and U7ex51-treated mice. Results are expressed as copies per µl of input RNA and shown as mean ± SEM. Data normality was assessed prior to statistical testing; one-way ANOVA test was applied when assumptions were met, otherwise a non-parametric Kruskal–Wallis test was used. \**P* < 0.05; \*\**P* < 0.01; \*\*\**P* < 0.001.

We further projected all subclustered immune cells spatially in samples from *mdx52*, ASO-treated, and U7ex51-treated groups (Fig. 6D). Cluster 1, identified as activated microglial cells, was specifically localized to the fiber tract and hypothalamic regions in ASO-treated samples, consistent with the spatial expression patterns of *Spp1*, *Cd68*, and *Trem2* (Fig. 6E). To determine whether this immune response is specific to the ASO chemistry used in this study (tcDNA), we further analyzed the expression of immune markers in additional *mdx52* samples treated with ASOs modified with the 2′-O-methoxyethylribose (MOE) chemistry by Droplet digital PCR (ddPCR) (Fig. 6F-G; Fig. S10). We observed similarly elevated expression of immune markers *Trem2* and *Aif1,* and also increased mRNA level of Tnf-α and *Cxcl10* in the MOE-treated samples, suggesting that the distinct immune dynamics may be associated with ASO treatment in general, rather than being specific to a particular chemical backbone.

## DISCUSSION

This study provides a detailed spatial map of *Dmd* isoform expression in the central nervous system (CNS) of the mouse brain, alongside an evaluation of exon skipping therapies using ASO and U7ex51 approaches. Our findings reveal distinct patterns of isoform localization and coexpression, as well as treatment-induced responses to exon 51 skipping, including cellular restoration of dystrophin isoforms and changes in immune cell dynamics under the ASO treatment.

Identifying multiple dystrophin isoforms in the brain remains challenging. Although recent advances have introduced *in situ* hybridization and sequencing-based approaches, both methods face significant limitations: *in situ* hybridization is constrained by the multiplexing capacity, while sequencing techniques are restricted by the long *Dmd* transcript lengths and low abundance in the brain, making accurate isoform mapping difficult. In this study, we provide the first detailed spatial mapping of multiple dystrophin isoforms in the mouse brain across annotated brain regions and cell types at single-cell resolution using Xenium.

We observed two distinct spatial expression patterns between the full-length *Dmd* isoforms and shorter isoforms. In the hippocampus, full-length isoforms were primarily expressed in IT-ET Glut cells of the CA region, while shorter isoforms were enriched in the granule neurons in the dentate gyrus, consistent with previous finding (*41*). Interestingly, we observed distinct expression pattern across isocortex layers: full-length *Dmd* isoforms were expressed at higher levels in pyramidal neurons located in deeper cortical layers, while shorter isoforms were enriched in GABAergic interneurons and astrocytes in layer 1, suggesting that *Dmd* isoform expression is cell-type and cortical layer specific. Previous studies have demonstrated that full-length dystrophin isoforms play an essential role in regulating gamma-aminobutyric acid type A receptors (GABA_A_Rs) organization and inhibitory synaptic transmission in the central nervous system (*42, 43*). In the sensorimotor cortex of *mdx* mice, loss of Dp427 leads to a redistribution of GABA_A_Rs subunits, with increased α1 and decreased α2 expression as well as a higher proportion of small α5-containing clusters (*44*). α5-containing GABA_A_ARs are indicated to mediate tonic inhibition in layer 5 pyramidal neurons of the rat cortex (*45, 46*). These results suggest that loss of Dp427 might disrupt the tonic inhibition of deep cortical layer pyramidal neurons, which may contribute to the motor and cognitive abnormalities observed in dystrophinopathies. Further investigation into the full-length isoforms and association with GABA_A_Rs subunits in deeper cortical layers could provide critical insights into the molecular mechanisms in DMD.

Albeit the overall distinct expression patterns of full-length and shorter *Dmd* isoforms, we observed multiple neuronal cell types coexpressing these isoforms, particularly one third of *Dmd* positive CTX-CGE GABA neurons showed coexpression of long and short isoforms. The coexpression of isoforms in these neurons might suggest a potential interplay or regulation. Previous studies have reported rare instances of *Dmd* isoform coexpression in neurons (*14*). Our results indicate more frequent coexpression events in specific neuronal cell types, supporting the notion of a cell-specific transcriptional program rather than local proximity or cellular neighborhood.

Our targeted spatial transcriptomics approach allowed the evaluation of two distinct exon skipping therapies targeting *Dmd* exon 51. Comparison of neonatal U7ex51 delivery with adult ASO administration highlights several important trends. Both approaches successfully induced exon 51 skipping in the brain; however, U7ex51 delivered at postnatal day 1 consistently achieved higher skipping efficiency and broader restoration across full-length *Dmd* isoforms, whereas ASO delivered at postnatal day 49 primarily restored only Dp427c. This difference is likely attributable to the earlier intervention window as well as the persistence and accumulation of antisense sequences produced from the AAV genome. Although these preliminary data are based on a limited number of animals, they provide encouraging evidence that neonatal delivery of U7ex51 may offer more comprehensive and durable dystrophin rescue in the CNS than adult ASO treatment. Future work with adequately powered cohorts will be particularly valuable to determine whether the enhanced molecular restoration observed after neonatal U7ex51 delivery translates into improved behavioral outcomes, potentially exceeding the partial behavioral rescue previously achieved with adult ASO administration (*47*).

While this difference could be entirely due to earlier treatment, the lack of Dp427m restoration in ASO-treated mice could potentially lead to the persistence of a vascular phenotype, which would otherwise be corrected by the U7ex51 approach. Interestingly, although the CA1 region is enriched in full-length *Dmd* isoforms, it exhibited relatively low levels of exon 51 skipping. This contrasts with previous findings from the same experimental setup, which reported approximately 20% exon skipping after seven weeks of ASO, possibly due to technical variability, differences in the tissue capture regions, or the limited sample size (n=3) in this study.

In our ASO-treated groups, we observed activation of a distinct microglial population in the brain. Follow-up analyses in 2′MOE-injected animals indicated that this response was not specific to a given chemical modification (tcDNA vs. MOE), suggesting that microglial activation may represent a general response to ASO administration rather than to a particular ASO backbone. This observation is consistent with previous reports of microglial activation following CNS delivery of 2′-O-methyl ASOs (*48*) and other ASO therapies (*49*), which collectively point to a broader, transient immune reaction elicited by ASO-based therapies in the brain. Such effects should be carefully considered in the design and development of future CNS-targeted treatments. In contrast, U7ex51-treated samples did not display obvious signs of microglial activation, suggesting a potential advantage of this vectorized approach; however, we cannot exclude that the earlier timing of U7ex51 administration also contributed to this difference.

Several limitations remain in our analysis. First, we had to exclude the Dp140 isoform because its signal was very low, likely due to reliance on a single probeset necessitated by the limited sequence space available in its 5′ UTR (Data file S1). It would be valuable to include Dp140 in further studies, given its association with impairments in verbal memory, attention, and executive function (*13, 50*). Recent studies have also shown that loss of Dp140 is linked to ASD-like behaviors and enhanced anxiety in the *mdx52* mice (*51, 52*). Second, the mouse cerebellum was not included in this study, despite its implication in DMD (*53, 54*) and the reported expression of multiple *Dmd* isoforms (*14, 55*). Moreover, the current Xenium panel allows spatial analysis of limited gene panels. Future developments will likely expand the number of genes included and introduce disease-specific panels, enabling more comprehensive and targeted investigations.

In summary, we provide a detailed map of cell-type and region specificity of dystrophin isoforms in the mouse brain and investigate how these patterns differ in the *mdx52* mice lacking the full-length and Dp140 isoforms. Additionally, we evaluated the efficacy of two exon 51 skipping therapies in restoring dystrophin expression in the brain. Our findings demonstrate the power of high-resolution spatial transcriptomics to resolve isoform-level complexity in situ, supporting its broader application in studying *Dmd* transcript diversity in the brain and guiding the development of future therapeutic strategies.

## MATERIALS AND METHODS

### Study design

The primary objective of this study was to characterize the spatial distribution of *Dmd* isoforms across distinct brain regions and cell types, and to assess exon 51 skipping efficacy following two therapeutic interventions. Mouse selection and sample numbers are detailed in Fig. 1A. Tissue sections were chosen based on anatomical regions with reported *Dmd* expression, ensuring relevance to the biological question. Given the absence of prior studies applying spatial transcriptomics at the isoform level, no power analysis was performed. All samples and cells passing Xenium platform quality control were retained for downstream analysis. Although wildtype sample #3 exhibited reduced signal intensity (Fig. S1-3), its transcriptomic profile remained consistent with known regional biology (Fig. S4-6), and was therefore included in the final dataset.

### Animals, injections and tissue collection

The *mdx52* mouse model,carrying a deletion of exon 52 in the *Dmd* gene, was developed by Dr. Katsuki Motoya’s team. This model was created by replacing exon 52 with a neomycin resistance gene, resulting in the absence of Dp427, Dp260, and Dp140 dystrophins, while maintaining the expression of Dp116, Dp71 and Dp40. The strain was backcrossed with C57BL/6J mice for over eight generations to ensure genetic stability. This mouse line was generously shared by Prof. Toshikuni Sasaoka (Department of Comparative & Experimental Medicine/Brain Research Institute, Niigata University, Japan). Breeding pairs were provided to our facility based at UVSQ in Montigny-le-Bretonneux (France), by Dr. Jun Tanihata and Dr. Shin’ichi Takeda (National Center of Neurology and Psychiatry, Tokyo, Japan). Heterozygous females were bred with *mdx52* males to produce *mdx52* males and their C57BL/6J wild-type (WT) littermates.

Genotyping was performed using PCR analysis of tail DNA. All animal care and experimental protocols adhered to European regulations (CEE 86/609/EEC and EU Directive 2010/63/EU) as well as French national guidelines (87/848) and were approved by the Paris Center-South Ethics Committee (no. 59).

Three adult *mdx52* mice (P49) were deeply anesthetized with a single intraperitoneal injection of ketamine (95 mg/kg) and medetomidine (1 mg/kg) and received bilateral intracerebroventricular (ICV) injections of tricyclo-DNA (tcDNA) antisense oligonucleotide (ASO) targeting *Dmd* exon 51 (400 µg total; 5 µl per hemisphere, infused at 0.5 µl/min) (*35*). Age-matched *mdx52* and WT littermates received PBS under identical conditions as controls. To investigate whether earlier intervention might yield higher dystrophin rescue, a separate experimental group of three neonatal *mdx52* mice (postnatal day 1, P1) were anesthetized by hypothermia and received bilateral ICV of a self-complementary AAV9 vector encoding a modified U7snRNA targeting exon 51 (scAAV9-U7-51M; 2 µl per hemisphere, 7 × 10¹¹ vg total) (*56*). Stereotaxic coordinates for adult ICV injections were –0.5 mm from bregma, 1 mm lateral, and –2 mm from dura, whereas for neonatal ICV injections coordinates were positioned at equal distance between lambda and bregma, 0.8–1 mm lateral to the sagittal suture, and 3 mm deep (*35, 57*). Littermates were housed in groups of two to five per cage under a 12-hour light/dark cycle (lights on at 7:00 a.m.) with *ad libitum* access to food and water. All mice were euthanized by cervical dislocation at 14 weeks of age. Brains were collected and processed as follows: one hemisphere was immediately frozen on dry ice for Xenium analysis, while the other hemisphere was dissected to isolate specific brain regions, including the hippocampus (HIP) and cortex (CX). These tissue samples were snap-frozen in liquid nitrogen and stored at –80 °C until further processing for RT-qPCR, ddPCR, and western blot experiments.

### Xenium sample preparation

Fresh-frozen hemi-brain samples were equilibrated to cryostat temperatures (−20 °C) for 30 minutes before mounting with Tissue-Tek® O.C.T. Compound (Sakura Finetek, Torrance, CA, USA) using fast freezing. Samples were cryo-sectioned at 10 µm using Leica CM3050 S cryostat (Leica Microsystems, Wetzlar, Germany) in coronal orientation and placed onto Xenium slides with randomization of placement, followed by fixation and permeabilization. Experiments were performed according to the manufacturer’s instructions (10x Genomics, CG000579 & CG000581). A total of 247 genes from the Xenium pre-designed brain panel and 50 genes from a custom panel were then applied to the slides (Data File. S1). All probes hybridized to their target RNAs, were ligated to form circular DNA, and enzymatically amplified to generate multiple copies of gene-specific barcodes for each transcript. Background autofluorescence was quenched, and nuclei were counterstained with DAPI, following the manufacturer’s protocol (10x Genomics, CG000582). Processed Xenium slides were loaded into the Xenium analyzer (v1.8.2.1).

### Xenium data analysis

Raw data were processed using Xenium Ranger (v1.7.1). Cell segmentation was based on DAPI-stained nuclei with a 15 μm expansion. Low-quality transcript signals (q < 20) were excluded from downstream analysis. Cells were assigned manually to corresponding samples using Xenium Explorer (v3.0).

Xenium data were then annotated at two levels: brain region and cell type. For brain region annotation, we used BANKSY v0.1.6 (*58*) with lamda = 0.8 to group the cells on the first Xenium slide into different brain regions. These regions were then visually matched to the anatomy map from the Allen Brain Atlas (*59*) based on the closest coronal section. Then, we used scANVI with scvi-tools v1.1.4 (*60*) to transfer the labels from the first slide to all other three slides that were not yet annotated. For cell type annotation, we used single-cell transcriptomics data (v2 chemistry) from the whole adult mouse brain from the Allen Brain Cell Atlas as a reference (*36*). First, we selected data from brain regions that matched those in our Xenium data, as identified in the region annotation described earlier. Then, we removed cell types with low expression levels and downsampled the remaining data to 10% for each cell type to reduce computational load. Finally, we transferred the cell type labels from the reference dataset to our Xenium data using scANVI (Fig. S11; Data file S2-3).

### Differential proportion analysis

We applied Scanpro v0.3.2 to compare cell type proportions between groups (*61*). First, we calculated the proportion of each cell type for every replicate within each group. Then, Scanpro was used to fit a linear model on these proportions across groups, using empirical Bayes shrinkage to stabilize the variance estimates. To account for the heteroskedasticity that can occur with proportions from a binomial distribution, Scanpro applied a logit transformation to all percentage values before testing, then using an empirical Bayes moderated t-test when comparing two groups. Finally, we adjusted the resulting *P*-values for multiple testing using the Benjamini–Hochberg (BH) method.

### Isoform localization and coexpression analysis

To calculate the (co)expression rates of the *Dmd* full-length and shorter isoforms across brain region and cell type pairs several steps were undertaken. Firstly, to remove brain regions with very few or no cells of that type, we kept only regions where each cell type made up more than 0.5% of the cells based on the cell type proportion map across regions (Fig. S4A). Secondly, we removed any region–cell type pairs where the isoform had fewer than 10 raw counts. Finally, for each region and cell type, we calculated the expression rate as the proportion of cells positive for each isoform, and the coexpression rate as the number of cells positive for both isoforms divided by the total number of cells positive for either isoform (i.e. the Jaccard index between cells expressing either one or both isoforms).

### Skipping rate calculation

For each region and cell type in every sample, we calculated the exon 51 skipping rate as:

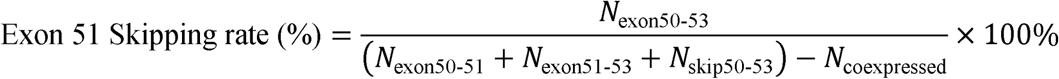

where N_exon50-51_ is the number of cells with unskipped probes, N_exon51-53_ is the number of cells of exon 52 deletion probes, N_skip50-53_ is the number of cells with exon 51 skipping probes, then we excluded N_coexpressed_ with the number of cells expressing multiple probes to avoid double counting. If no cells were detected in these categories, the skipping rate was set to zero.

### Differential expression analysis between exon 51 positive and negative cells

To address the large imbalance in cell numbers between the exon 51-skipped positive and negative populations during differential expression analysis, we designed a permutation test. From the exon 51-skipped negative population, we selected a subset of cells that matched the positive population in terms of region and cell type distribution. If the pool of matching cells was not enough to match the number of exon 51-skipped positive population, we expanded the selection pool further based on matching the cell type distribution alone. If that pool was also exhausted, we further relaxed the criteria to match only by region. In the ASO group, each cell in the exon 51-skipped negative population could be selected up to five times, whereas in the U7ex51 group, each cell could be selected up to seven times, due to the higher numbers of exon 51-positive cells and the need to maintain balanced sampling between positive and negative populations. We applied sc.tl.rank_genes_groups() from Scanpy using the Wilcoxon test, with Benjamini-Hochberg correction for multiple testing. The permutation test was run 1,000 times. For each gene, we recorded the number of times it showed a significant difference and calculated the average log-fold change across all iterations as the effect size.

### Western blot

Protein extracts were prepared from brain tissues using RIPA lysis and extraction buffers (Thermo Fisher Scientific, Waltham, MA, USA), supplemented with SDS to a final concentration of 5% (Bio-Rad, France). Protein concentrations were measured with the BCA Protein Assay Kit (Thermo Fisher Scientific, Waltham, MA, USA). Samples were denatured at 100 °C for 3 min, and 25 µg of protein per sample was loaded onto NuPAGE 3–8% Tris-acetate gels (Invitrogen), according to the manufacturer’s protocol. Dystrophin expression was analyzed using the anti-dystrophin monoclonal rabbit antibody (ab154168, Abcam, France), while vinculin served as a loading control and was detected with the hVin-1 monoclonal mouse antibody (V9131, Sigma, Saint-Louis, USA). Detection was performed with an IRDye 800CW goat anti-mouse secondary antibody and an IRDye 680CW goat anti-rabbit secondary antibody (Li-Cor, Germany), and bands were visualized using the Odyssey CLx imaging system (Li-Cor). Quantification was carried out with Image Studio software (Li-Cor), normalizing dystrophin signals to vinculin levels. Standard curves were generated for each brain region by mixing lysates from wild-type (WT) and *mdx52* control samples to obtain defined dystrophin percentages (0%, 2.5%, 5%, 10%, and 20% relative to WT tissues).

### RNA analyses

Total RNA was isolated from dissected brain structures using TRIzol reagent according to the manufacturer’s instructions (Thermo Fisher Scientific). For visualization of exon-skipping efficacy on gels, 1 µg of total RNA was subjected to RT-PCR analysis using the LunaScript RT SuperMix Kit (New England Biolabs) in a 20 μL reaction. cDNA synthesis was carried out at 55 °C for 10 min. PCR was then performed from 1.5 µL of cDNA with GoTaq G2 Colorless Master Mix (Promega) in a 25 µL reaction, using primers Ex-m50F (5′-AGGAAGTTAGAAGATCTGAGG-3′) and Ex-m55R (5′-GGAACTGCTGCAGTAATCTATGA-3′) under the following cycling conditions: 32 cycles of 95 °C for 30 s, 58 °C for 1 min, and 72 °C for 1 min. PCR products were electrophoresed on 1.5% agarose gels and quantified with ImageLab Software (Bio-Rad). Exon 51 skipping was also quantified by TaqMan real-time PCR as previously described (*62*), using specific assays targeting the exon 50–51 junction (assay Mm.PT.58.41685801; forward primer: 5′-CAAAGCAGCCTGACCGT-3′; reverse primer: 5′-TGACAGTTTCCTTAGTAACCACAG-3′; probe: 5′-TGGACTGAGCACTACTGGAGCCT-3′) and the exon 50–53 junction (forward primer: 5′-GCACTACTGGAGCCTTTGAA-3′; reverse primer: 5′-TTCCAGCCATTGTGTTGAATC-3′; probe: 5′-ACAGCTGCAGAACAGGAGACAACA-3′) (Integrated DNA Technologies). For each reaction, 150 ng of cDNA was used, and all measurements were performed in triplicate. qPCR was conducted under fast cycling conditions using a Bio-Rad CFX384 Touch Real-Time PCR Detection System. Data analysis was based on the absolute quantification method: copy numbers of the exon-skipped (exon 50–53) and unskipped (exon 50–51) products were determined using gBlock standards Ex49–54Δ52 and Ex49–54Δ51+52 (Integrated DNA Technologies). Exon 51 skipping levels were expressed as the percentage of total dystrophin transcripts, calculated as the ratio of skipped copies to the sum of skipped and unskipped copies.

Droplet digital PCR (ddPCR) was performed on the same RNA samples used for RT-PCR and qPCR analyses to quantify the expression of inflammatory markers. After reverse transcription, cDNA was subjected to ddPCR using the QX200 Droplet Digital PCR system (Bio-Rad), according to the manufacturer’s instructions. Specific assays were used to detect the following transcripts: TREM2 (assay Mm.PT.58.7992121), AIF1 (forward primer: 5′-CCCACCGTGTGACATCCA-3′; reverse primer: 5′-TGGTCCCCCAGCCAAGA-3′; probe: 5′-AGCTATCTCCGAGCTGCCCTGATTGG-3′), LAPTM5 (assay Mm.PT.58.31533533), TNF-α (forward primer: 5′-TGGGAGTAGACAAGGTACAACCC-3′; reverse primer: 5′-CATCTTCTCAAAATTCGAGTGACAA-3′; probe: 5′-CACGTCGTAGCAAACCACCAAGTGGA-3′), TGF-β1 (assay Mm.PT.58.11254750), CXCL10 (assay Mm.PT.58.43575827) (Integrated DNA Technologies), CDKN1A (assay Mm01303209_m1) and IL6 (assay Mm00446190_m1) (Thermo Fisher Scientific).

Each ddPCR reaction was prepared in a final volume of 40 µL containing 50 ng of cDNA, ddPCR Supermix for Probes (Bio-Rad), and the corresponding primer/probe assay. Droplets were generated with the QX200 Droplet Generator (Bio-Rad), and PCR amplification was performed on a C1000 Touch Thermal Cycler (Bio-Rad) under the following conditions: 95 °C for 10 min, 39 cycles of 94 °C for 30 s and 55 °C for 1 min, and a final step at 98 °C for 10 min. Droplets were then read on a QX200 Droplet Reader (Bio-Rad), and absolute transcript quantification was obtained using QuantaSoft Analysis Pro software (Bio-Rad). Results were expressed as copies per µL of input RNA. No-template controls were included in each run to monitor potential contamination.

### Statistical analysis

Statistical analyses were conducted using R v4.3.2, Python v3.10, and GraphPad Prism v10.3.3. Group comparisons were performed using linear mixed model regression with wildtype as the reference. For comparisons involving more than two groups, post-hoc pairwise analyses were carried out using estimated marginal means (emmeans) with Tukey’s adjustment for multiple testing. Comparison of *Dmd* isoforms and regional probes between wildtype and *mdx52* mice were assessed by the linear regression test with Benjamini–Hochberg correction. Futher, comparison of selected *Dmd* isoforms and regional across brain regions and cell types were assessed using one-sided t-tests with Benjamini–Hochberg correction. Non-parametric Mann– Whitney U tests were applied to evaluate qPCR and Western blot results. For ddPCR results, data normality was assessed prior to statistical testing, one-way ANOVA test was applied when assumptions were met, otherwise a non-parametric Kruskal–Wallis test was used. A significance threshold of α = 0.05 was used throughout, statistical significance was defined as \**P* < 0.05; \*\**P* < 0.01; \*\*\**P* < 0.001.

## List of Supplementary Materials

Fig S1 to S11 for supplementary figures Data files S1 to S3

## Acknowledgments

We acknowledge funding by the Medical Genomics Theme and the Human Genetics department of the LUMC. We thank Dr. Katsuki Motoya, Prof. Sasaoka Toshikuni (Department of Comparative & Experimental Medicine/ Brain Research Institute; Niigata University, Japan), Dr. Jun Tanihata and Dr. Shin’ichi Takeda (National Center of Neurology and Psychiatry, Tokyo, Japan) for providing the mdx52 mouse breeders. We also thank Sandra de Haan (Leiden University Medical Center; Leiden, The Netherlands) for reviewing the manuscript and Lola van Doorne for performing the Xenium experiment.

## Author contributions

Each author’s contribution(s) to the paper should be listed [we encourage you to follow the CRediT model]. Each CRediT role should have its own line, and there should not be any punctuation in the initials.

Examples:

Conceptualization: QM, AA

Methodology: QM, AA, OV, MD, AG, AM, PS

Investigation: SdV, LH, OV, MD

Visualization: QM, AA, OV, MD

Funding acquisition: PS, AG, MVP

Project administration: PS, AM

Supervision: PS, AM, MVP, and AAR

Writing – original draft: QM

Writing – review & editing: all authors

## Data and materials availability

All data and materials used in this study are publicly available. Processed Xenium sequencing data, including raw expression matrices and annotated metadata can be accessed through [https://doi.org/10.5281/zenodo.17285218]. All analysis scripts and code used for data preprocessing and visualization are provided in a public GitHub repository (https://github.com/Qirongmao97/Xenium_DMD_brain).As reference of cell type annotation, adult mouse scRNA-seq (v2 chemistry) from the Allen Brain Atlas can be downloaded in the public data repository (https://allen-brain-cell-atlas.s3.us-west-2.amazonaws.com/index.html#expression_matrices/WMB-10Xv2/20230630/)

## Supplementary Materials

### Data files

**Data file. S1:** List of genes and *Dmd* probes included in the Xenium panel

**Data file. S2:** Marker gene list for cell type annotation in Xenium data

**Data file. S3:** Marker gene list for regional annotation in Xenium data

## Supplementary Figures

**Fig. S1.**
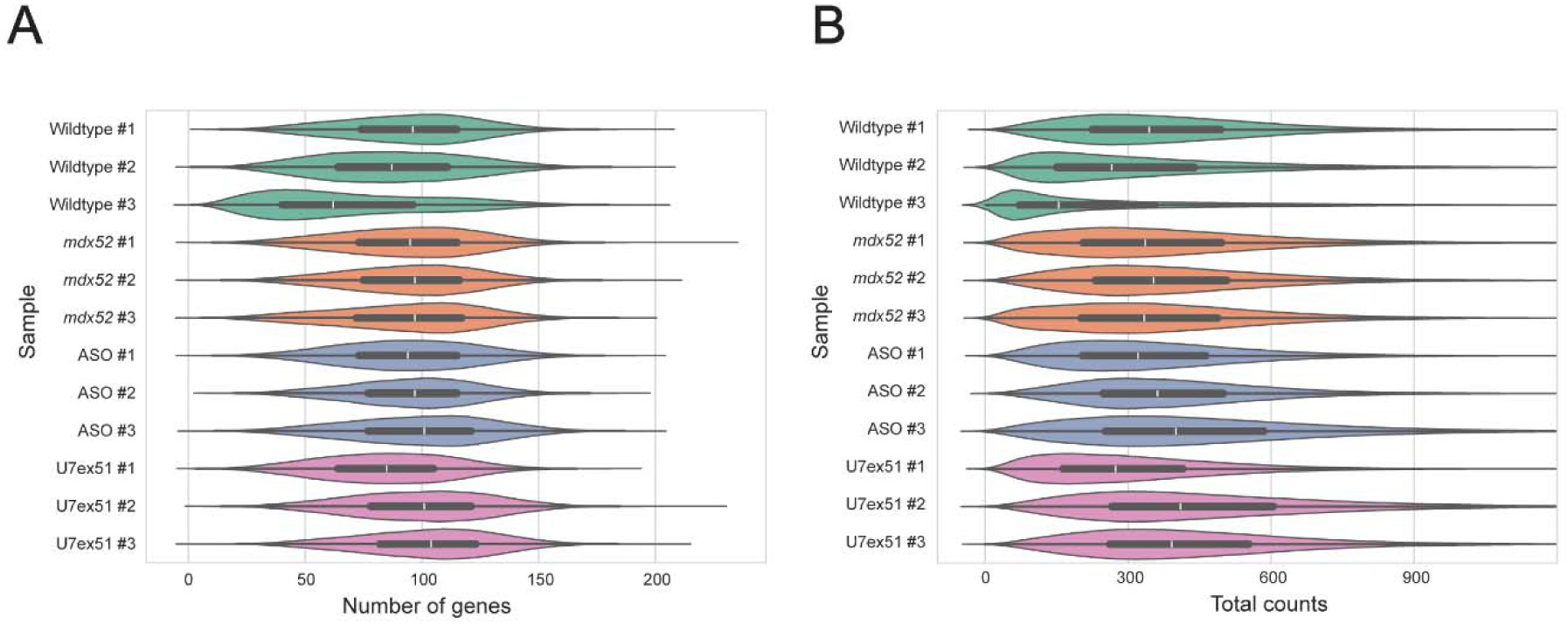
Quality view of Xenium data. Violin plots showing **(A)** the number of genes detected per cell and **(B)** the total counts per cell across samples and experimental groups.

**Fig. S2.**
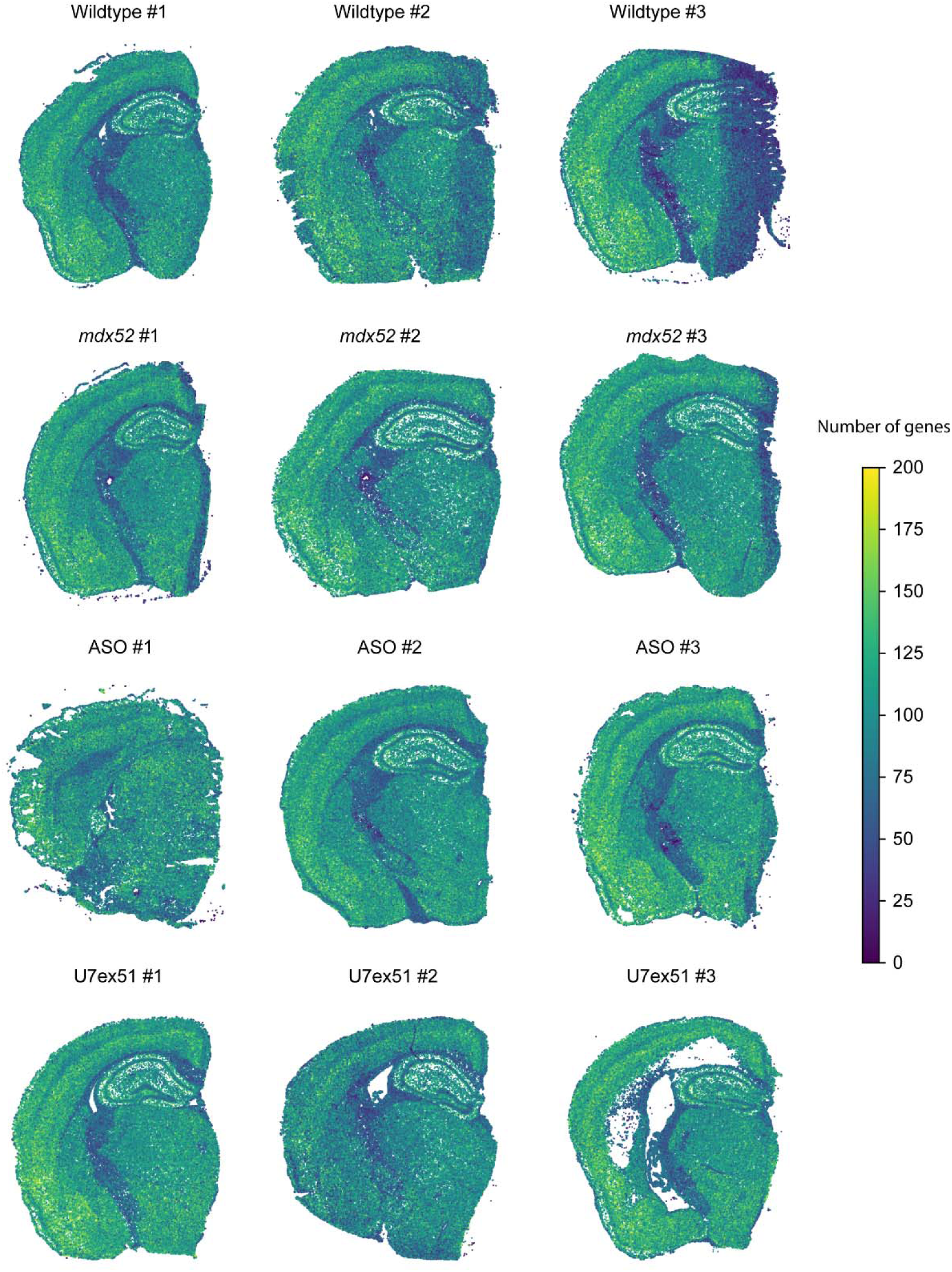
Spatial distribution of number of genes across samples. Spatial plot of number of genes distribution across samples.

**Fig. S3.**
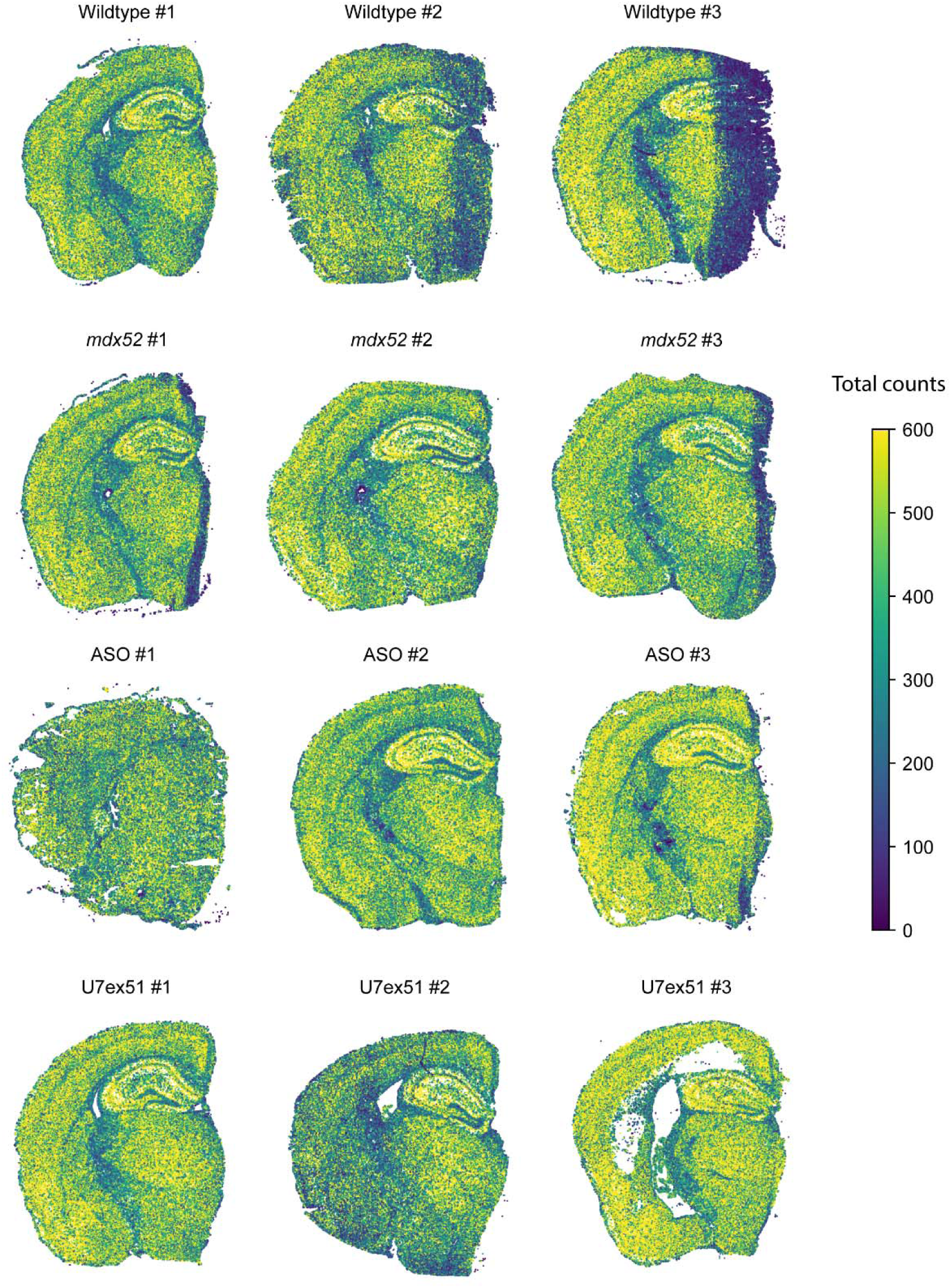
Spatial distribution of total counts across samples. Spatial plot of total counts distribution across samples

**Fig. S4.**
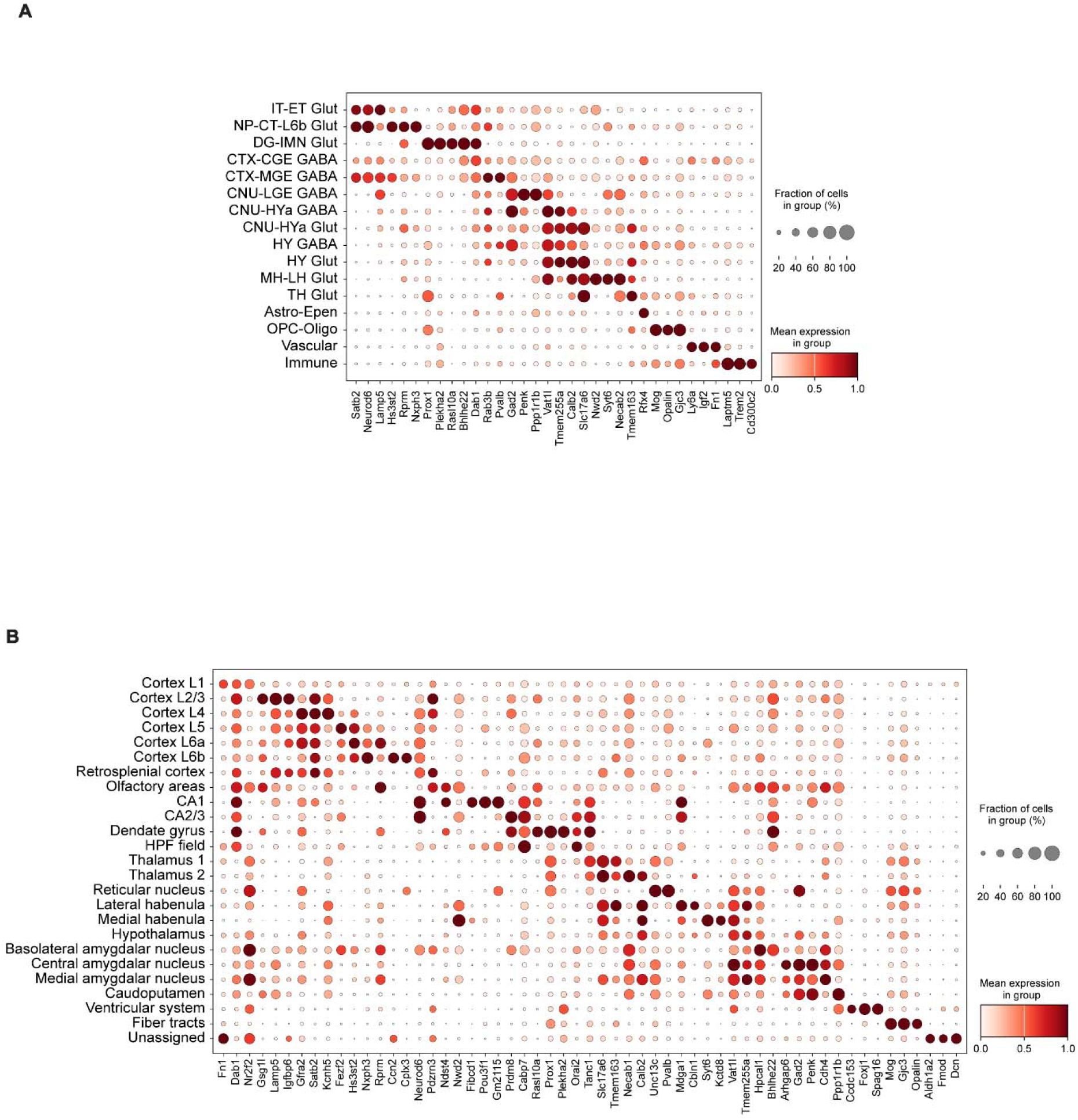
Marker genes of annotated clusters. Dot plots showing the mean expression of the top three marker genes for each annotated **(A)** cell type and **(B)** brain region group.

**Fig. S5.**
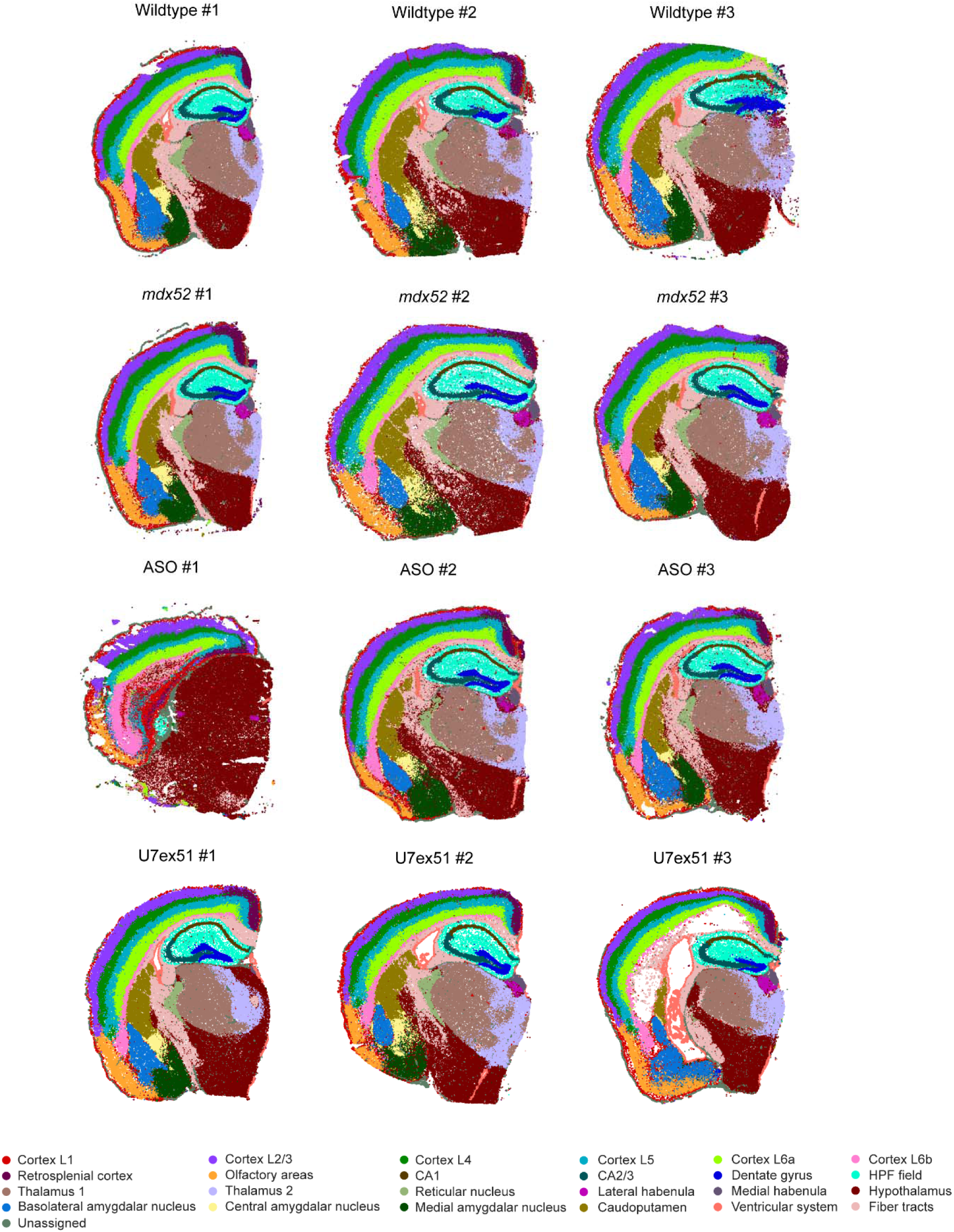
Regional annotation for all samples. Spatial plot of regional annotations across samples.

**Fig. S6.**
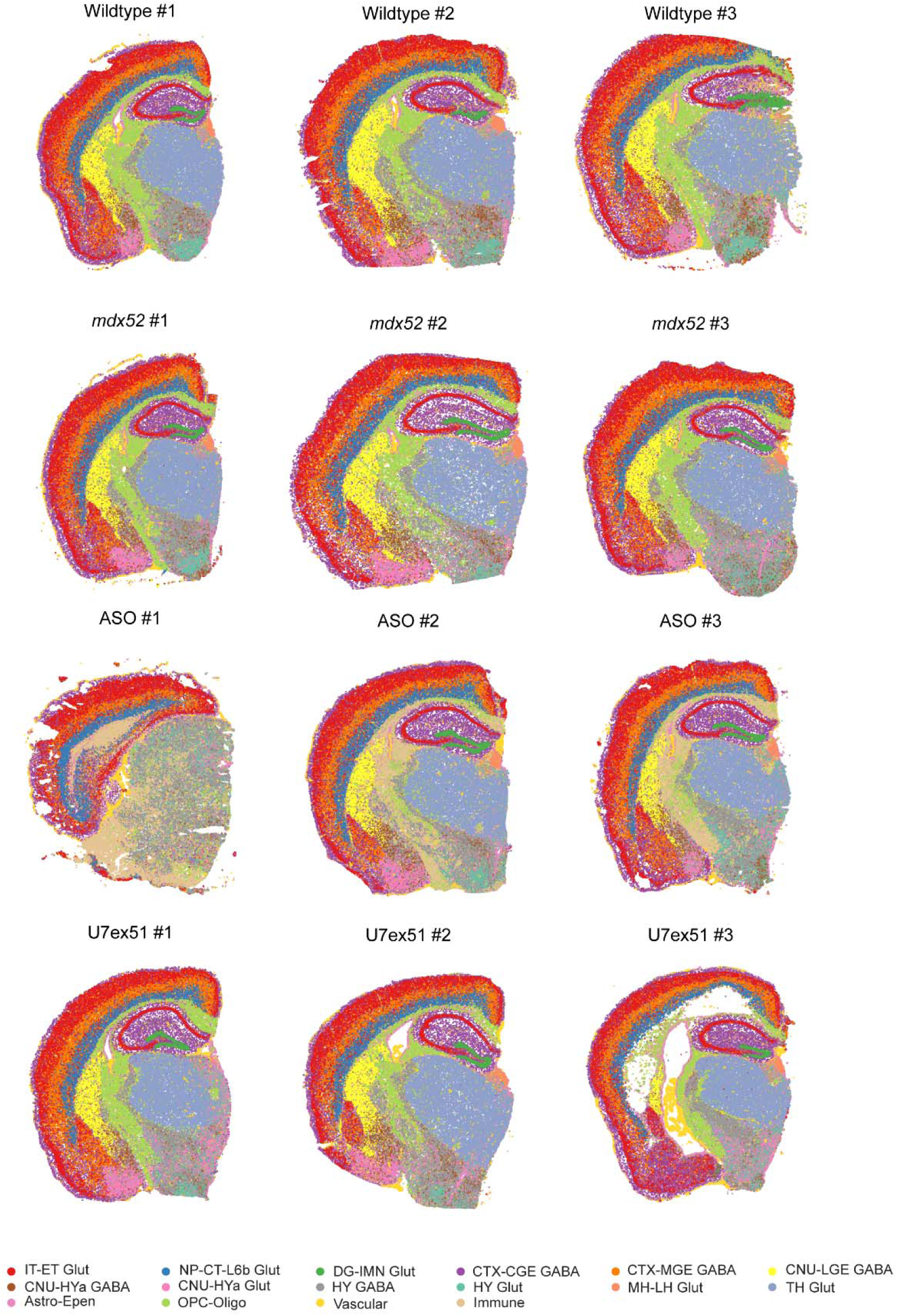
Cell type annotation for all samples. Spatial plot of cell type annotations across samples.

**Fig. S7.**
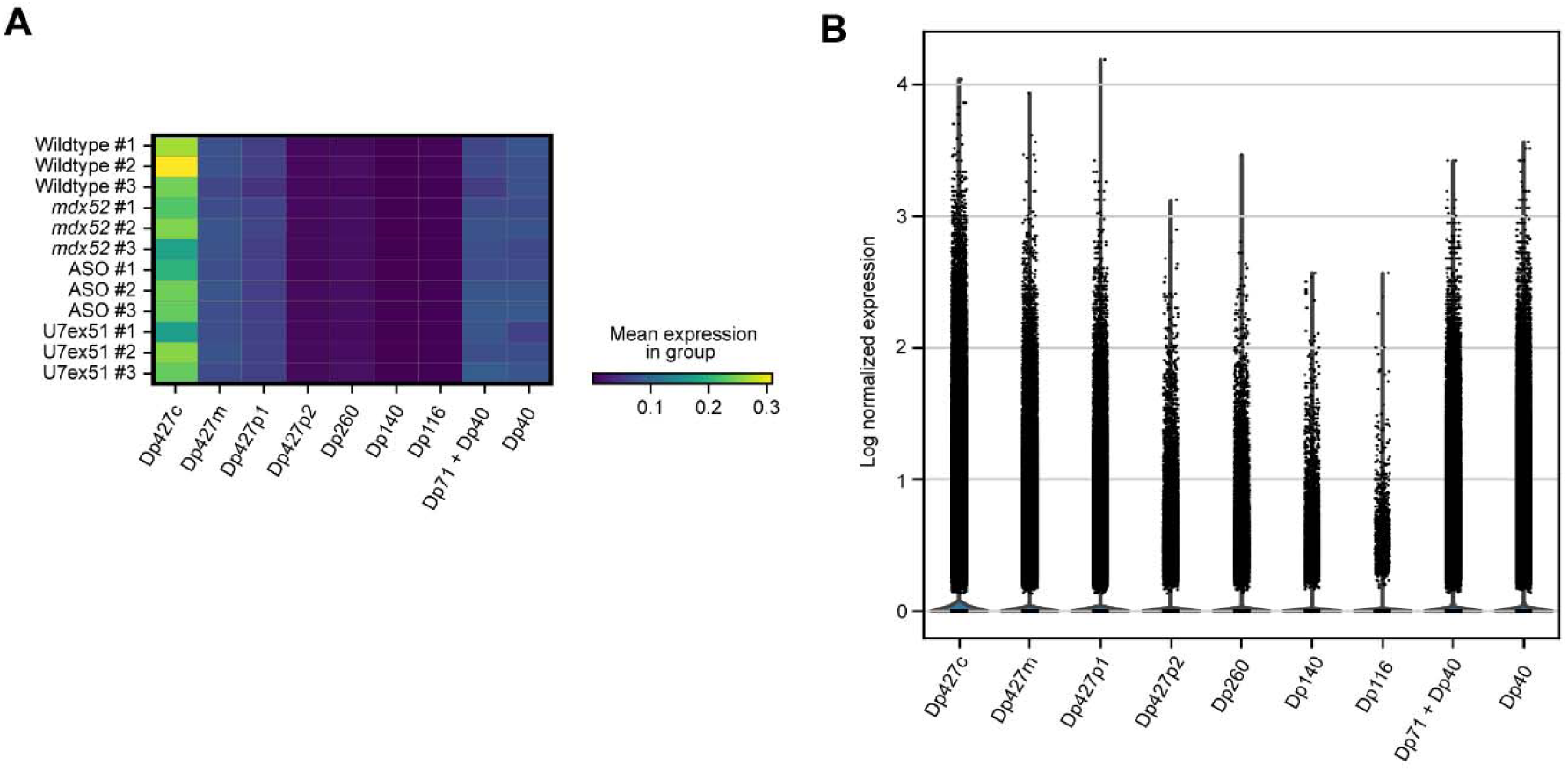
Quality of Xenium custom probes targeting *Dmd* isoforms. **(A)** Heatmap showing log-normalized mean expression levels of *Dmd* isoforms across samples targeted by Xenium custom probes. **(B)** Violin plot illustrating overall *Dmd* isoform expression across all samples; each dot represents expression in an individual cell.

**Fig. S8.**
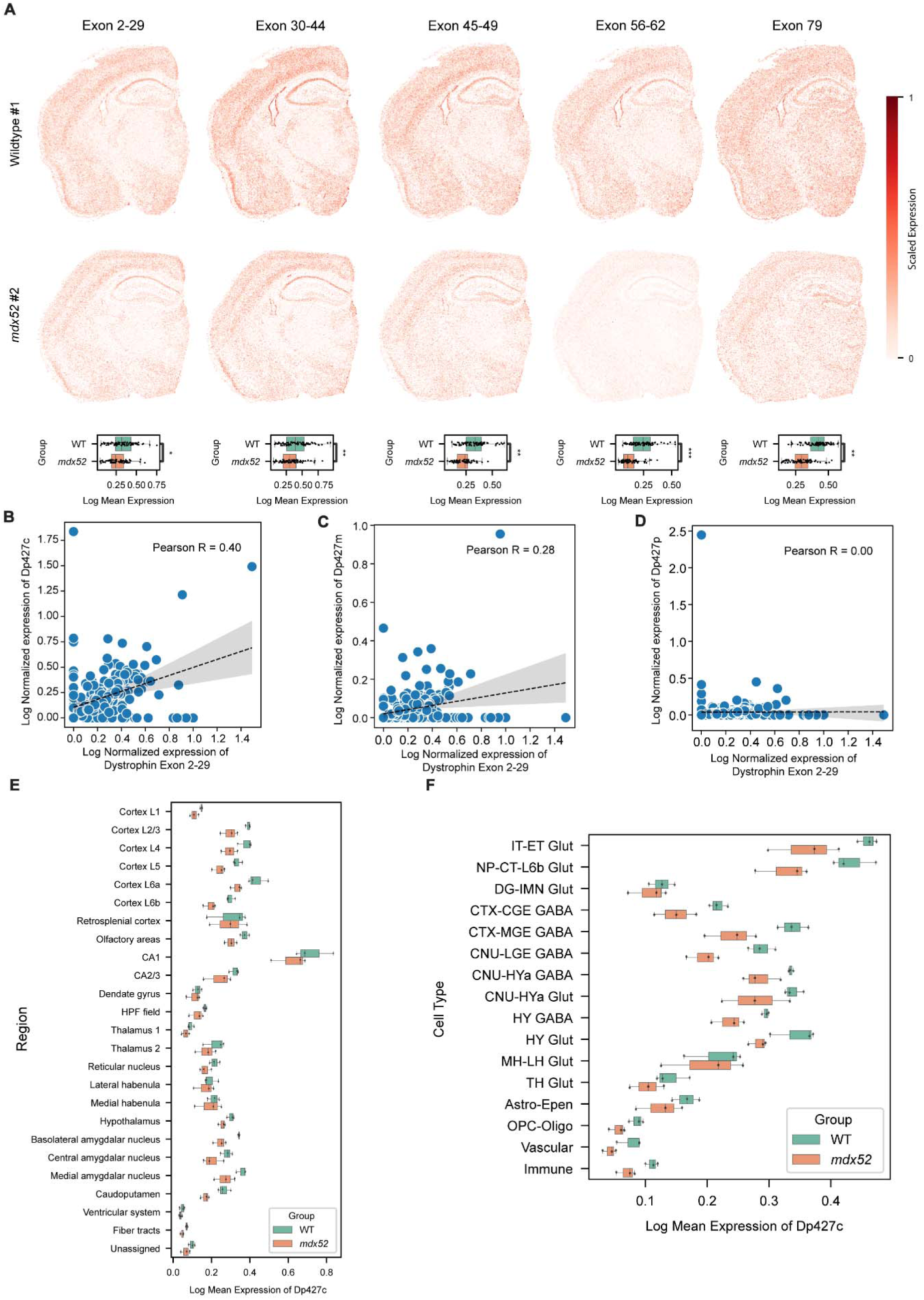
Regional probe expression across wildtype and *mdx52* samples. **(A)** (Top) Spatial distribution of *Dmd* regional probes in WT and *mdx52* mouse brains colored by scale expression of each *Dmd* regional probe. (Bottom) Boxplot with overlaid dotplot showing the log-transformed mean expression of each *Dmd* regional probe across brain regions per sample across different groups. Statistical analysis by the linear regression test with BH correction. **(B-D)** Dot plot showing the correlation between *Dmd* regional probe exon 2-29 and **(B)** Dp427c; **(C)** Dp427m; **(D)** Dp427p. **(E-F)** Expression of Dp427c across brain regions and cell types in WT and *mdx52* mice. Statistical analysis by the one-sided t-test with BH correction. \**P* < 0.05; \*\**P* < 0.01; \*\*\**P* < 0.001.

**Fig. S9.**
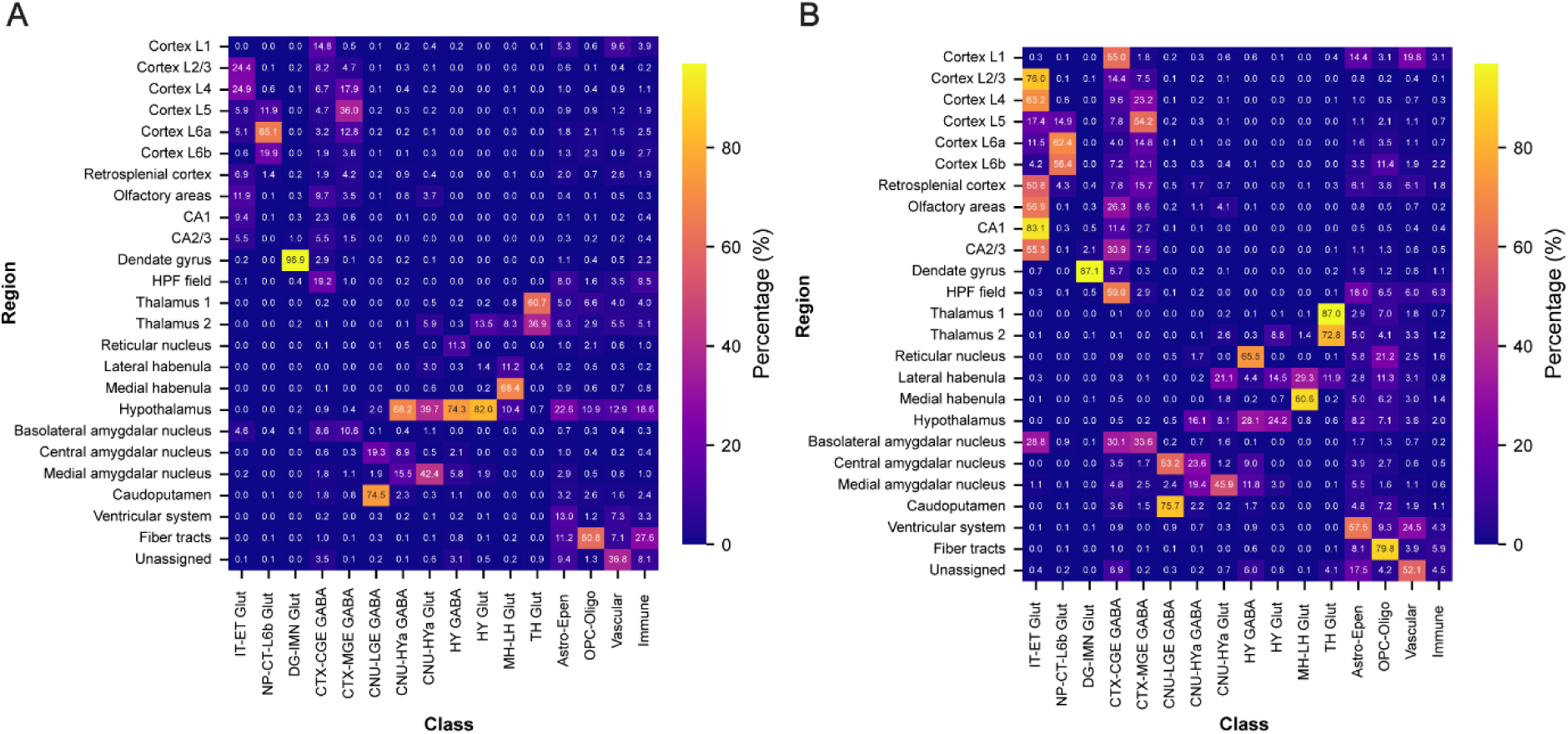
Cell type composition and distribution across regions. Each row corresponds to a brain region and each column to a cell type. (**A**) Distribution of cell types across regions: For each cell type, the heatmap shows the fraction of its total cells located in each region, highlighting how each cell type is spatially distributed (column sums = 100%). (**B**) Cell type composition per region: For each region, the heatmap shows the proportion of each cell type within that region, illustrating the relative cellular makeup of each region (row sums = 100%).

**Fig. S10.**
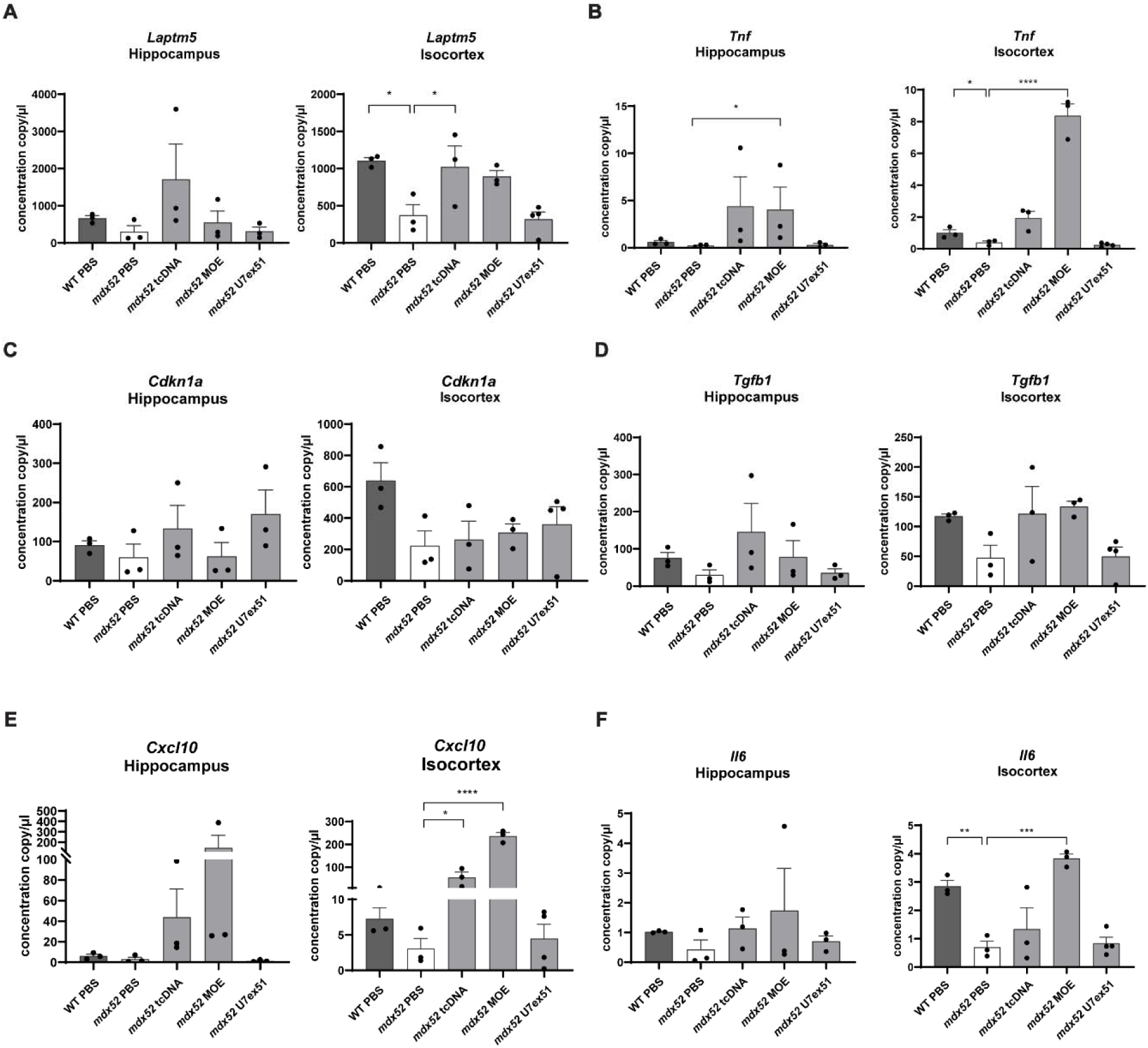
Quantification of immune marker expression in brains treated with ASOs of different chemical backbones. **(A–F)** Absolute quantification of mRNA levels measured by droplet digital PCR (ddPCR) in the hippocampus (left) and isocortex (right) for: **(A)** *Laptm5* (\**p* = 0.0197 WT vs *mdx52*; \**p* = 0.0379 *mdx52* PBS vs tcDNA51), **(B)** *Tnf* (hippocampus \**p* = 0.0412; isocortex \**p* = 0.0472), **(C)** *Cdkn1a*, **(D)** *Tgfb1*, **(E)** *Cxcl10* (\**p* = 0.0283; \*\*\*\**p* < 0.0001), and **(F)** *Il6* (\**p* = 0.0067; \*\**p* = 0.0004). Experimental groups include wild type mice control (WT PBS), *mdx52* mice control (*mdx52 PBS*), *mdx52* tcDNA51, *mdx52* MOE51, and *mdx52* AAV-U7-51. Results are expressed as copies per µL of input RNA and shown as mean ± SEM (n = 3–4 per group). Data normality was assessed prior to statistical testing; one-way ANOVA test was applied when assumptions were met, otherwise a non-parametric Kruskal–Wallis test was used. Statistical significance between groups is indicated by horizontal bars.

**Fig. S11.**
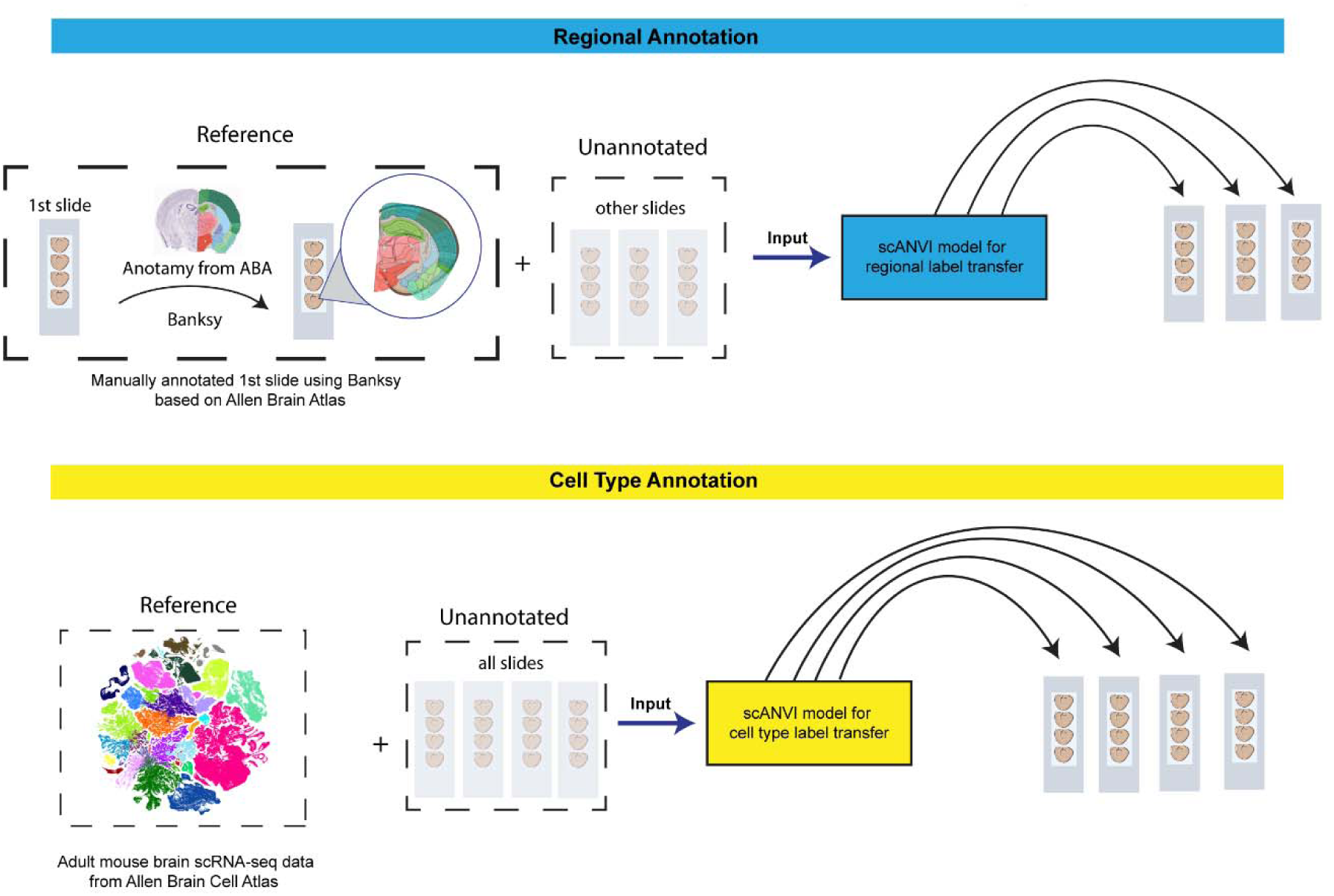
Schematic diagram of annotation strategy of Xenium data. Schematic diagram of Xenium data annotation: (Top) Regional annotation: the first Xenium slide was annotated using Banksy, guided by anatomical structures from the Allen Brain Atlas. This annotated slide, along with other unannotated slides, was used as input to the scANVI model for label transfer. (Bottom) Cell type annotation: A well-annotated, downsampled scRNA-seq atlas of the adult mouse brain from the Allen Brain Atlas was used as a reference. All unannotated slides were used as input to the scANVI model for cell type label transfer.

## References and Notes

1. Cotton S, Voudouris NJ, Greenwood KM. Intelligence and Duchenne muscular dystrophy: full-scale, verbal, and performance intelligence quotients. Dev Med Child Neurol. 2001;43(7):497–501.

2. Weerkamp PMM, Mol EM, Sweere DJJ, Schrans DGM, Vermeulen RJ, Klinkenberg S, et al. Wechsler Scale Intelligence Testing in Males with Dystrophinopathies: A Review and Meta-Analysis. Brain Sci. 2022;12(11).

3. Gregg J, Wilson C, Curran D, Hanna D. Neurocognitive functioning among children and young people with Duchenne Muscular Dystrophy: A systematic review and meta-analysis. Clin Neuropsychol. 2024;38(8):1806–33.

4. Darmahkasih AJ, Rybalsky I, Tian CX, Shellenbarger KC, Horn PS, Lambert JT, et al. Neurodevelopmental, behavioral, and emotional symptoms common in Duchenne muscular dystrophy. Muscle Nerve. 2020;61(4):466–74.

5. Pane M, Lombardo ME, Alfieri P, D’Amico A, Bianco F, Vasco G, et al. Attention Deficit Hyperactivity Disorder and Cognitive Function in Duchenne Muscular Dystrophy: Phenotype-Genotype Correlation. J Pediatr-Us. 2012;161(4):705-+.

6. Wu JY, Kuban KCK, Allred E, Shapiro F, Darras BT. Association of Duchenne muscular dystrophy with autism spectrum disorder. J Child Neurol. 2005;20(10):790–5.

7. Lee AJ, Buckingham ET, Kauer AJ, Mathews KD. Descriptive Phenotype of Obsessive Compulsive Symptoms in Males With Duchenne Muscular Dystrophy. J Child Neurol. 2018;33(9):572–9.

8. Monaco AP, Neve RL, Colletti-Feener C, Bertelson CJ, Kurnit DM, Kunkel LM. Isolation of candidate cDNAs for portions of the Duchenne muscular dystrophy gene. Nature. 1986;323(6089):646-50.

9. Pane M, Messina S, Bruno C, D’Amico A, Villanova M, Brancalion B, et al. Duchenne muscular dystrophy and epilepsy. Neuromuscular Disord. 2013;23(4):313–5.

10. Billard C, Gillet P, Barthez MA, Hommet C, Bertrand P. Reading ability and processing in Duchenne muscular dystrophy and spinal muscular atrophy. Developmental Medicine and Child Neurology. 1998;40(1):12–20.

11. Astrea G, Pecini C, Gasperini F, Brisca G, Scutifero M, Bruno C, et al. Reading impairment in Duchenne muscular dystrophy: A pilot study to investigate similarities and differences with developmental dyslexia. Res Dev Disabil. 2015;45–46:168-77.

12. Hinton VJ, De Vivo DC, Fee R, Goldstein E, Stern Y. Investigation of Poor Academic Achievement in Children with Duchenne Muscular Dystrophy. Learn Disabil Res Pract. 2004;19(3):146–54.

13. Doorenweerd N, Mahfouz A, van Putten M, Kaliyaperumal R, t’ Hoen PAC, Hendriksen JGM, et al. Timing and localization of human dystrophin isoform expression provide insights into the cognitive phenotype of Duchenne muscular dystrophy (vol 7, 12575, 2017). Sci Rep-Uk. 2018;8.

14. Catapano F, Alkharji R, Chambers D, Singh S, Aghaeipour A, Malhotra J, et al. A comprehensive spatiotemporal map of dystrophin isoform expression in the developing and adult human brain. Acta Neuropathol Commun. 2025;13(1):110.

15. Dsouza VN, Man NT, Morris GE, Karges W, Pillers DAM, Ray PN. A Novel Dystrophin Isoform Is Required for Normal Retinal Electrophysiology. Human Molecular Genetics. 1995;4(5):837–42.

16. Byers TJ, Lidov HGW, Kunkel LM. An Alternative Dystrophin Transcript Specific to Peripheral-Nerve. Nature Genetics. 1993;4(1):77–81.

17. Tetorou K, Aghaeipour A, Singh S, Morgan JE, Muntoni F. The role of dystrophin isoforms and interactors in the brain. Brain. 2025;148(4):1081–98.

18. Pascual-Morena C, Cavero-Redondo I, Sequí-Domínguez I, Rodríguez-Gutiérrez E, Visier-Alfonso ME, Martínez-Vizcaíno V. Intelligence quotient-genotype association in dystrophinopathies: A systematic review and meta-analysis. Neuropath Appl Neuro. 2023;49(3).

19. Pascual-Morena C, Cavero-Redondo I, Martínez-Vizcaíno V, Sequí-Domínguez I, Fernández-Bravo-Rodrigo J, Jiménez-López E. Dystrophin Genotype and Risk of Neuropsychiatric Disorders in Dystrophinopathies: A Systematic Review and Meta-Analysis. J Neuromuscular Dis. 2023;10(2):159–72.

20. Ricotti V, Mandy WPL, Scoto M, Pane M, Deconinck N, Messina S, et al. Neurodevelopmental, emotional, and behavioural problems in Duchenne muscular dystrophy in relation to underlying dystrophin gene mutations. Developmental Medicine and Child Neurology. 2016;58(1):77–84.

21. Chamova T, Guergueltcheva V, Raycheva M, Todorov T, Genova J, Bichev S, et al. Association between loss of dp140 and cognitive impairment in duchenne and becker dystrophies. Balk J Med Genet. 2013;16(1):21–9.

22. Chesshyre M, Ridout D, Hashimoto Y, Ookubo Y, Torelli S, Maresh K, et al. Investigating the role of dystrophin isoform deficiency in motor function in Duchenne muscular dystrophy. J Cachexia Sarcopeni. 2022;13(2):1360–72.

23. Muntoni F, Signorovitch J, Sajeev G, Lane H, Jenkins M, Dieye I, et al. DMD Genotypes and Motor Function in Duchenne Muscular Dystrophy: A Multi-institution Meta-analysis With Implications for Clinical Trials. Neurology. 2023;100(15):e1540–e54.

24. Aartsma-Rus A, Fokkema I, Verschuuren J, Ginjaar L, van Deutekom J, van Ommen GJ, et al. Theoretic Applicability of Antisense-Mediated Exon Skipping for Duchenne Muscular Dystrophy Mutations. Hum Mutat. 2009;30(3):293–9.

25. Heo YA. Golodirsen: First Approval. Drugs. 2020;80(3):329–33.

26. Shirley M. Casimersen: First Approval. Drugs. 2021;81(7):875–9.

27. Mendell JR, Rodino-Klapac LR, Sahenk Z, Roush K, Bird L, Lowes LP, et al. Eteplirsen for the Treatment of Duchenne Muscular Dystrophy. Ann Neurol. 2013;74(5):637–47.

28. Hoy SM. Delandistrogene Moxeparvovec: First Approval. Drugs. 2023;83(14):1323–9.

29. Gadgil A, Raczynska KD. U7 snRNA: A tool for gene therapy. J Gene Med. 2021;23(4).

30. Schümperli D, Pillai R. The special Sm core structure of the U7 snRNP:: far-reaching significance of a small nuclear ribonucleoprotein. Cell Mol Life Sci. 2004;61(19-20):2560–70.

31. Goyenvalle A, Vulin A, Fougerousse F, Leturcq F, Kaplan JC, Garcia L, et al. Rescue of dystrophic muscle through U7 snRNA-mediated exon skipping. Science. 2004;306(5702):1796-9.

32. Vulin A, Barthélémy I, Goyenvalle A, Thibaud JL, Beley C, Griffith G, et al. Muscle Function Recovery in Golden Retriever Muscular Dystrophy After AAV1-U7 Exon Skipping. Mol Ther. 2012;20(11):2120–33.

33. Phase I/IIa Systemic Gene Delivery Clinical Trial of scAAV9.U7.ACCA for Exon 2 Duplication-Associated Duchenne Muscular Dystrophy [Internet]. 2020. Available from: https://clinicaltrials.gov/study/NCT04240314.

34. Doorenweerd N, Mahfouz A, van Putten M, Kaliyaperumal R, t’ Hoen PAC, Hendriksen JGM, et al. Timing and localization of human dystrophin isoform expression provide insights into the cognitive phenotype of Duchenne muscular dystrophy. Sci Rep-Uk. 2017;7.

35. Saoudi A, Barberat S, le Coz O, Vacca O, Doisy Caquant M, Tensorer T, et al. Partial restoration of brain dystrophin by tricyclo-DNA antisense oligonucleotides alleviates emotional deficits in mdx52 mice. Mol Ther Nucleic Acids. 2023;32:173-88.

36. Yao ZZ, van Velthoven CTJ, Kunst M, Zhang M, Mcmillen D, Lee CKY, et al. A high-resolution transcriptomic and spatial atlas of cell types in the whole mouse brain. Nature. 2023;624(7991).

37. Goyenvalle A, Jimenez-Mallebrera C, van Roon W, Sewing S, Krieg AM, Arechavala-Gomeza V, et al. Considerations in the Preclinical Assessment of the Safety of Antisense Oligonucleotides. Nucleic Acid Ther. 2023;33(1):1–16.

38. Schmid CD, Sautkulis LN, Danielson PE, Cooper J, Hasel KW, Hilbush BS, et al. Heterogeneous expression of the triggering receptor expressed on myeloid cells-2 on adult murine microglia. J Neurochem. 2002;83(6):1309–20.

39. Ladwig A, Walter HL, Hucklenbroich J, Willuweit A, Langen KJ, Fink GR, et al. Osteopontin Augments M2 Microglia Response and Separates M1-and M2-Polarized Microglial Activation in Permanent Focal Cerebral Ischemia. Mediat Inflamm. 2017;2017.

40. Perego C, Fumagalli S, De Simoni MG. Temporal pattern of expression and colocalization of microglia/macrophage phenotype markers following brain ischemic injury in mice. J Neuroinflamm. 2011;8.

41. Fujimoto T, Yaoi T, Nakano K, Arai T, Okamura T, Itoh K. Generation of dystrophin short product-specific tag-insertion mouse: distinct Dp71 glycoprotein complexes at inhibitory postsynapse and glia limitans. Cell Mol Life Sci. 2022;79(2).

42. Suzuki Y, Higuchi S, Aida I, Nakajima T, Nakada T. Abnormal distribution of GABA(A) receptors in brain of duchenne muscular dystrophy patients. Muscle Nerve. 2017;55(4):591–5.

43. Knuesel I, Mastrocola M, Zuellig RA, Bornhauser B, Schaub MC, Fritschy JM. Short communication: altered synaptic clustering of GABAA receptors in mice lacking dystrophin (mdx mice). Eur J Neurosci. 1999;11(12):4457–62.

44. Zarrouki F, Goutal S, Vacca O, Garcia L, Tournier N, Goyenvalle A, et al. Abnormal Expression of Synaptic and Extrasynaptic GABA(A) Receptor Subunits in the Dystrophin-Deficient mdx Mouse. Int J Mol Sci. 2022;23(20).

45. Jang HJ, Cho KH, Kim MJ, Yoon SH, Rhie DJ. Layer-and cell-type-specific tonic GABAergic inhibition of pyramidal neurons in the rat visual cortex. Pflugers Arch. 2013;465(12):1797–810.

46. Yamada J, Furukawa T, Ueno S, Yamamoto S, Fukuda A. Molecular basis for the GABAA receptor-mediated tonic inhibition in rat somatosensory cortex. Cereb Cortex. 2007;17(8):1782–7.

47. Saoudi A, Barberat S, le Coz O, Caquant MD, Tensorer T, Sliwinski E, et al. Partial restoration of brain dystrophin by tricyclo-DNA antisense oligonucleotides alleviates emotional deficits in mice. Mol Ther Nucl Acids. 2023;32:173–88.

48. Toonen LJA, Casaca-Carreira J, Pellisé-Tintoré M, Mei HL, Temel Y, Jahanshahi A, et al. Intracerebroventricular Administration of a 2′-O-Methyl Phosphorothioate Antisense Oligonucleotide Results in Activation of the Innate Immune System in Mouse Brain. Nucleic Acid Ther. 2018;28(2):63–73.

49. Amanat M, Nemeth CL, Fine AS, Leung DG, Fatemi A. Antisense Oligonucleotide Therapy for the Nervous System: From Bench to Bedside with Emphasis on Pediatric Neurology. Pharmaceutics. 2022;14(11).

50. Tyagi R, Aggarwal P, Mohanty M, Dutt V, Anand A. Computational cognitive modeling and validation of Dp140 induced alteration of working memory in Duchenne Muscular Dystrophy. Sci Rep-Uk. 2020;10(1).

51. Hashimoto Y, Kuniishi H, Sakai K, Fukushima Y, Du X, Yamashiro K, et al. Brain Dp140 alters glutamatergic transmission and social behaviour in the mdx52 mouse model of Duchenne muscular dystrophy. Prog Neurobiol. 2022;216:102288.

52. Saoudi A, Zarrouki F, Sebrie C, Izabelle C, Goyenvalle A, Vaillend C. Emotional behavior and brain anatomy of the mdx52 mouse model of Duchenne muscular dystrophy. Dis Model Mech. 2021;14(9).

53. Cyrulnik SE, Hinton VJ. Duchenne muscular dystrophy: a cerebellar disorder? Neurosci Biobehav Rev. 2008;32(3):486–96.

54. Vicari S, Piccini G, Mercuri E, Battini R, Chieffo D, Bulgheroni S, et al. Implicit learning deficit in children with Duchenne muscular dystrophy: Evidence for a cerebellar cognitive impairment? PLoS One. 2018;13(1):e0191164.

55. Hendriksen RG, Schipper S, Hoogland G, Schijns OE, Dings JT, Aalbers MW, et al. Dystrophin Distribution and Expression in Human and Experimental Temporal Lobe Epilepsy. Front Cell Neurosci. 2016;10:174.

56. Vacca O, Saoudi A, Doisy M, Phongsavanh X, Le Coz O, Nagy C, et al. Ineffective behavioral rescue despite partial brain Dp427 restoration by AAV9-U7-mediated exon 51 skipping in *mdx52* mice. bioRxiv. 2025:2025.09.19.677185.

57. Kim JY, Grunke SD, Levites Y, Golde TE, Jankowsky JL. Intracerebroventricular Viral Injection of the Neonatal Mouse Brain for Persistent and Widespread Neuronal Transduction. Jove-J Vis Exp. 2014(91).

58. Singhal V, Chou NG, Lee JS, Yue YF, Liu JY, Chock WK, et al. BANKSY unifies cell typing and tissue domain segmentation for scalable spatial omics data analysis. Nature Genetics. 2024;56(3).

59. Wang Q, Ding SL, Li Y, Royall J, Feng D, Lesnar P, et al. The Allen Mouse Brain Common Coordinate Framework: A 3D Reference Atlas. Cell. 2020;181(4):936–53 e20.

60. Xu CL, Lopez R, Mehlman E, Regier J, Jordan M, Yosef N. Probabilistic harmonization and annotation of single-cell transcriptomics data with deep generative models. Mol Syst Biol. 2021;17(1).

61. Alayoubi Y, Bentsen M, Looso M. Scanpro is a tool for robust proportion analysis of single-cell resolution data. Sci Rep-Uk. 2024;14(1).

62. Aupy P, Zarrouki F, Sandro Q, Gastaldi C, Buclez PO, Mamchaoui K, et al. Long-Term Efficacy of AAV9-U7snRNA-Mediated Exon 51 Skipping in mdx52 Mice. Mol Ther-Meth Clin D. 2020;17:1037–47.

